# Probiotic-Directed Fermentation Reprograms the Metabolic Profile of a Traditional Mongolian Whole-Wheat Diet and Modulates Escherichia coli-Induced Gut Microbiota Dysbiosis

**DOI:** 10.64898/2026.08.08.743650

**Authors:** Eerdun duleng, Qingchun Ling, Jinhua Bao, Surigaga, Tegexi, Nandien, Shandan, Aruhan, Yuhong Bai, A Lisha, Cuiqin Gong, Burenbatu, Shengliang Ni, Wenping

## Abstract

Traditional Mongolian fermented foods have been extensively utilized for dietary regulation and the promotion of gastrointestinal health. However, spontaneous fermentation remains inherently unpredictable, leading to significant variations in microbial community dynamics, metabolite accumulation, and the consistency and quality of the final product. Drawing on the traditional preparation of Mongolian acidic foods, this study established a controlled production strategy for whole-wheat probiotic fermented soup (WWPFS) by combining enzymatic pretreatment with probiotic-directed fermentation.

Physicochemical characterization, 16S rRNA gene-based microbial community profiling, LC–MS/MS-based untargeted metabolomics, safety evaluation, and an Escherichia coli-induced gut microbiota dysbiosis model were employed to optimize and comprehensively characterize the fermentation process of WWPFS. The optimized process established a reproducible fermentation system consistently dominated by Lactobacillus and Bacillus across independent fermentation batches. Compared with traditional spontaneous fermentation, probiotic-directed fermentation remodeled the physicochemical properties of the whole-wheat matrix, including carbon, nitrogen, phosphorus, sulfur, and mineral composition, and facilitated the accumulation of putatively annotated LC–MS/MS features, including DL-lactate, 1,4-D-xylobiose, diacetyl, and phenyllactic-acid-related features derivatives. Acute oral and 28-day repeated-dose toxicity evaluations showed no treatment-related adverse effects within the tested dose range and study duration. In the Escherichia coli-induced gut microbiota dysbiosis mouse model, microbial richness, diversity, and community structure differed among the experimental groups, and both low- and high-dose WWPFS groups showed significant shifts in overall gut microbial community composition relative to the model group after multiple-testing correction, together with directional recovery of selected model-responsive bacterial genera. Cross-system integration identified coordinated response patterns between fermentation-derived metabolite features and model-responsive gut bacterial taxa, supporting a potential metabolite–microbiota link in WWPFS-mediated gut microbiota modulation. In summary, probiotic-directed fermentation improved the controllability of the traditional Mongolian fermented food production process, reshaped its metabolic profile, and enhanced its potential to modulate the gut microbiota. These findings provide experimental evidence supporting the modernization of traditional Mongolian fermented foods and the development of probiotic-based functional foods.

## 1. Introduction

Traditional Mongolian fermented foods represent an integral component of Mongolian medicine and reflect the longstanding principle of food–medicine homology. These products are generally produced through the microbial transformation of grains, plant-derived materials, dairy products, or mixed substrates, and have historically been used for dietary regulation, gastrointestinal support, immune maintenance, and recovery from fatigue [1–3]. During fermentation, complex microbial communities can degrade macromolecular substrates and generate organic acids, oligosaccharides, peptides, phenolic acid derivatives, and other metabolites, which may collectively contribute to the physiological properties of the final product [4–6].

Although traditional fermented dietary preparations have substantial historical value, their reliance on spontaneous fermentation makes the process difficult to standardize. Fermentation outcomes are readily affected by environmental microorganisms, raw-material variability, temperature, fermentation time, humidity, and empirical processing conditions, resulting in inconsistent microbial composition, fluctuating metabolite profiles, unstable sensory and nutritional properties, and uncertain safety boundaries [7,8]. These limitations restrict quality control, scaled production, and evidence-based development of traditional fermented diets.

Probiotic-directed fermentation provides a feasible strategy to improve the controllability and functional consistency of traditional fermented products. By introducing selected strains with defined metabolic capacities, fermentation can be guided toward predictable substrate utilization, organic acid production, microbial stabilization, and formation of beneficial metabolites [4]. Previous studies have shown that probiotic fermentation can improve cereal substrate conversion, reduce anti-nutritional constraints, enhance phenolic release, and generate metabolites relevant to gastrointestinal health [7–13].

A traditional Mongolian acidic dietary preparation is recorded in the classical Mongolian medical text Four parts of Mongolian medicine nectar and represents a typical food–medicine fermented preparation [14]. In traditional preparation, fermented wheat mash is pressed to obtain the liquid fraction, followed by decoction with dried ginger, butter, and brown sugar. Historically, this preparation has been used for digestive discomfort, fatigue and dietary regulation, particularly among older adults [15,16]. However, its current preparation still largely depends on spontaneous fermentation and empirical processing, leading to complex composition, limited reproducibility, and potential safety concerns.

Therefore, this study aimed to establish a controllable WWPFS preparation system for this traditional Mongolian acidic dietary preparation using enzymatic pretreatment and probiotic- directed fermentation. Enzymatic hydrolysis and fermentation parameters were first optimized, followed by systematic evaluation of microbial community structure, physicochemical and nutritional features, and metabolite profiles. Safety tests and an E. coli-induced gut microbiota dysbiosis model were further used to assess its biological relevance.

## 2. Materials and Methods

### 2.1 Materials, Reagents, and Microbial Strains

Dried ginger ( No. 20220608), brown sugar (food grade), butter (food grade), cellulase ( No. C8270), pectinase ( No. P8181), reference standards, analytical reagents, digestive enzymes, buffers, and Hetao wheat from Inner Mongolia, China were used in this study. Detailed information on materials, reagents, standards, batch numbers, suppliers, and uses is provided in Supplementary Table S1. Freeze-dried powders of Lactobacillus plantarum (No. BNCC187897), Lactobacillus casei (No. BNCC 186562), and Bacillus subtilis were (No. BNCC19462) , Escherichia coli (No. BNCC 3336435) purchased from commercial suppliers. Unless otherwise specified, all chemicals were of analytical or chromatographic grade. Animal experiments were approved by the Medical Ethics Committee of the Affiliated Hospital of Inner Mongolia Minzu University (Approval No. NM-LL-2024-04-25-67; approval date: April 25, 2024) and conducted in accordance with institutional guidelines and relevant national regulations for laboratory animal welfare and ethical review.

### 2.2 Process Study of the WWPFS Fermentation System

#### 2.2.1 Preparation of the Enzymatic Hydrolysate

##### (1) Raw-material pretreatment and enzymatic hydrolysis

Whole-wheat grains and dried ginger were separately ground and passed through a 100-mesh sieve. Whole-wheat powder, ginger powder, brown sugar, and butter were mixed in equal proportions and suspended in purified water at a solid-to-liquid ratio of 1:5 (m/v). The pH was adjusted to 6.0 with 1 mol/L NaOH [17]. Cellulase and pectinase were then added according to the experimental design, and the mixture was incubated under the specified enzymatic hydrolysis conditions. The reaction was terminated by heating, and the hydrolysate was cooled for subsequent experiments.

##### (2) Single-factor experiments for enzymatic hydrolysis optimization

Single-factor experiments were performed to evaluate the effects of enzymatic hydrolysis conditions. The investigated factors included total enzyme dosage (0.1%, 0.2%, 0.3%, 0.4%, 0.5%, and 0.6%), cellulase-to-pectinase ratio (1:7, 3:5, 1:1, 5:3, and 7:1), hydrolysis time (10, 20, 30, 40, and 50 min), and hydrolysis temperature (17, 27, 37, 47, and 57 °C). Evaluation indices and parameter selection were based on previous studies [18–20]. Total flavonoid and total polyphenol contents were determined by spectrophotometry and calculated using the corresponding standard curves (Supplementary Fig. S1a-b). According to the single-factor results, hydrolysis time, hydrolysis temperature, and total enzyme dosage were selected for further optimization using an L9(3^4) orthogonal design. Detailed experimental layouts and results are provided in Supplementary Tables S2 and S3 and Supplementary Fig. S1c-f.

To minimize interference from other variables, the cellulase-to-pectinase ratio was fixed at 1:1, and the enzymatic hydrolysis pH was maintained at 6.0 in subsequent experiments. All measurements were performed in triplicate.

##### (3) Comprehensive evaluation of enzymatic hydrolysis efficiency

Hydrolysates obtained under different conditions were ultrasonicated at 40 °C and 50 Hz for 30 min, followed by centrifugation at 8000 rpm for 15 min. The supernatant was collected for analysis.

A multi-criteria decision-making (MCDM) method was used to comprehensively evaluate enzymatic hydrolysis efficiency [21]. The composite score (Y; full score, 100) was calculated from total flavonoid content (Y1) and total polyphenol content (Y2), with each assigned a weight of 50%, using the formula Y = (Y1/Y1max) × 50 + (Y2/Y2max) × 50. Detailed scoring definitions are provided in Supplementary Tables S2 and S3.

The optimal enzymatic hydrolysis conditions were selected based on the composite score, total flavonoid content, and total polyphenol content.

#### 2.2.2 Screening of Fermentation Strains

According to the International Scientific Association for Probiotics and Prebiotics (ISAPP) consensus statement and studies on cereal and plant-based fermentation [4,22,23], Lactobacillus plantarum, Bacillus subtilis, and Lactobacillus casei were selected as candidate strains.

The Oxford cup method was used to evaluate the adaptability and growth performance of different strains in individual substrates and enzymatic hydrolysate [24]. Whole-wheat powder, ginger powder, brown sugar, and enzymatic hydrolysate were used as culture substrates. Growth around the Oxford cups and inhibition zones were observed to determine growth-promoting or inhibitory effects [25,26].

The results showed that whole-wheat powder, ginger powder, brown sugar, and enzymatic hydrolysate supported the growth of *L. plantarum* and *B. subtilis*, whereas *L. casei* was inhibited to varying degrees. Therefore, *L. plantarum* and *B. subtilis* were selected as the probiotic consortium for fermentation. Detailed strain-screening results are provided in Supplementary Fig. S1g-i.

#### 2.2.3 Optimization of WWPFS Fermentation Conditions

The optimized enzymatic hydrolysate was sealed with a heat-resistant plant tissue culture sealing film (microporous membrane) and aluminum foil, sterilized at 121 °C for 15 min, and cooled to room temperature. The selected probiotic consortium was inoculated into the sterilized hydrolysate according to the experimental design and fermented under controlled conditions in a constant-temperature shaking incubator. This sealing approach allowed gas exchange while reducing dust and microbial contamination, representing a non-forced-aeration and relatively oxygen-limited fermentation condition. Samples were collected under the specified single-factor conditions for subsequent analysis.

##### (1) Single-factor experiments

Single-factor experiments were conducted to investigate the effects of fermentation conditions on WWPFS preparation. The evaluated factors included starter dosage (hereafter using inoculum size), with 6%, 8%, 10%, 12%, and 14% groups. *L. plantarum* and *B. subtilis* ratio (1:9, 3:7, 1:1, 7:3, and 9:1), fermentation time (12, 16, 20, 24, and 28 h), and fermentation temperature (27, 32, 37, 42, and 47 °C).

The results showed that fermentation time, fermentation temperature, and inoculum size significantly affected fermentation performance, whereas the strain ratio did not show a significant effect within the tested range. Therefore, the *L. plantarum* and *B. subtilis* ratio was fixed at 1:1 in subsequent experiments. Detailed single-factor fermentation results are provided in Supplementary Table S5 and Supplementary Fig. S1j-m.

##### (2) Response surface optimization

Based on the single-factor results, fermentation time, fermentation temperature, and inoculum size were selected as independent variables [27]. A three-factor, three-level Box-Behnken design was used for response surface optimization.

Total flavonoid content (Y1), total polyphenol content (Y2), and viable cell count (Y3) were selected as response variables, and fermentation performance was evaluated using a weighted composite score: Y = (Y1/Y1max) × 25 + (Y2/Y2max) × 25 + (Y3/Y3max) × 50.

In the above equations, Y1max, Y2max, and Y3max indicate the maximum observed values of the corresponding indices within each optimization experiment.

Design-Expert software was used for model construction, significance analysis, and prediction of optimal conditions. Validation experiments were conducted under the predicted optimal conditions to assess model reliability and reproducibility. The detailed response surface design and validation scheme are provided in Supplementary Table S6.

### 2.3 Preparation of the Traditional Fermentation Control

According to traditional whole-wheat fermented product preparation methods [14,15], fermented wheat mash was first pressed to obtain the liquid fraction, followed by addition of ginger powder, butter, and brown sugar and decoction at 75 °C for 25 min. The resulting product was used as the traditional fermentation control for comparison with WWPFS in terms of physicochemical characteristics, chemical composition, and metabolite profile.

### 2.4 Transformation Characteristics of Functional Components

To evaluate the transformation characteristics of functional components during enzymatic hydrolysis and fermentation, seven treatment groups were established: S1, raw-material group; S2, traditional fermentation group (CTDH); S3, enzymatic hydrolysate group; S4, heat-inactivated bacteria fermentation group; S5, *B. subtilis* fermentation group; S6, *L. plantarum* fermentation group; and S7, probiotic consortium fermentation group (DH). Three independent batches were prepared for each group.

pH was measured using a digital pH meter [28]. Total organic acids were determined by acid-base titration [29]. Soluble dietary fiber (SDF) was measured using the AOAC enzymatic-gravimetric method [30]. Total soluble solids (TSS) were determined by the constant-weight method [31]. Total carbon, total nitrogen (TN), non-protein nitrogen (NPN), total phosphorus (TP), total sulfur (TS), magnesium (Mg), and manganese (Mn) were measured according to national standard methods [32,33]. The NPN/TN ratio was calculated as the ratio of non-protein nitrogen to total nitrogen.

All measurements were performed in triplicate. Detailed analytical procedures and reagent information are provided in the Supplementary Materials and Supplementary Table S1.

### 2.5 Microbial Community Analysis of WWPFS Fermentation Products

Total microbial DNA was extracted from six independent WWPFS fermentation batches using a HiPure DNA extraction kit (Magen, Guangzhou, China). DNA extraction, PCR amplification, library preparation, sequencing, read processing, OTU construction, and primary taxonomic annotation were performed by Gene Denovo Biotechnology Co., Ltd. (Guangzhou, China). The V3-V4 region of the bacterial 16S rRNA gene was amplified using primers 341F (5′-CCTACGGGNGGCWGCAG-3′) and 806R (5′-GGACTACHVGGGTATCTAAT-3′). PCR was performed in triplicate under the following conditions: 94 °C for 2 min; 30 cycles of 98 °C for 10 s, 62 °C for 30 s, and 68 °C for 30 s; and a final extension at 68 °C for 5 min. Amplicons were purified using an AxyPrep DNA Gel Extraction Kit, quantified using an ABI StepOnePlus Real-Time PCR System, pooled in equimolar amounts, and sequenced using paired-end 250-bp reads on an Illumina HiSeq 2500 platform.

Raw reads containing more than 10% ambiguous nucleotides or fewer than 50% of bases with Q > 20 were removed using fastp. Paired-end reads were merged using FLASH v1.2.11 with a minimum overlap of 10 bp and a maximum mismatch rate of 2%. Merged tags were further filtered using QIIME v1.9.1, and chimeric sequences were removed using UCHIME against reference release r20110519. Effective tags were clustered into operational taxonomic units at 97% sequence similarity using UPARSE v9.2.64. The most abundant sequence within each OTU was selected as its representative sequence. Taxonomic assignments were performed using the RDP Classifier v2.2 against SILVA v132 with a confidence threshold of 0.8.

The resulting annotated OTU count tables were provided to the authors for downstream analysis. All diversity calculations, taxonomic aggregation, statistical testing, and figure generation reported in this study were performed independently by the authors in R 4.5.2.

For the six WWPFS fermentation batches, OTU counts were normalized to sample totals and expressed as relative abundance percentages. Genus-level profiles were obtained by summing the relative abundances of OTUs sharing the same provider-assigned genus annotation. OTUs without genus-level assignments were labelled according to the nearest classified parent taxon for visualization. Observed OTUs, Shannon diversity, Simpson diversity, and Pielou evenness were recalculated directly from the OTU count table. Because each fermentation batch contributed one microbial profile, these indices and genus-level abundances were summarized descriptively without batch-level inferential testing.

### 2.6 Untargeted Metabolomic Analysis

Six independent batches of traditional fermentation products and six independent batches of probiotic-fermentation products were prepared for untargeted metabolomic analysis. LC–MS/MS analysis was performed by Gene Denovo Biotechnology Co., Ltd. Metabolites were extracted using precooled methanol/acetonitrile/water (2:2:1, v/v/v), followed by low-temperature sonication for 30 min, incubation at −20 °C for 10 min, and centrifugation at 14,000 × g for 20 min at 4 °C. The supernatants were dried, reconstituted in 100 μL acetonitrile/water (1:1, v/v), and centrifuged at 14,000 × g for 15 min at 4 °C.

LC–MS/MS analysis was performed using an Agilent 1290 Infinity ultrahigh-performance liquid chromatography system coupled to an AB SCIEX TripleTOF 6600 mass spectrometer. Chromatographic separation was performed using an ACQUITY UPLC BEH Amide column (2.1 × 100 mm, 1.7 μm), with electrospray ionization operated in positive- and negative-ion modes. TOF-MS and product-ion spectra were acquired over m/z 60–1,000 and m/z 25–1,000, respectively, using information-dependent acquisition.

Raw files were converted to mzXML format using ProteoWizard and processed using XCMS with the centWave algorithm. Features detected in more than 50% of samples in at least one biological group were retained. Missing values were imputed using k-nearest-neighbour imputation, and feature intensities were normalized to the total peak area of each sample. LC– MS/MS features were putatively annotated by matching tandem mass spectra against MassBank, METLIN, MoNA, and an in-house spectral library. Because individual annotations were not confirmed using authentic standards and retention-time matching, all reported chemical identities were considered putative annotations.

### 2.7 Safety Evaluation

#### 2.7.1 Acute Oral Toxicity Test

The acute oral toxicity test was conducted according to the National Food Safety Standard: Acute Oral Toxicity Test (GB 15193.3-2014) [36,37].

Twenty SPF-grade ICR mice (8 weeks old; half male and half female) were randomly assigned to the administration group and control group (n = 10 per group).

The administration group received a single oral dose of WWPFS at 8.0 g/kg body weight, approximately equivalent to 40 times the recommended human intake. The control group received an equal volume of purified water (40 mL/kg body weight).

Animals were observed continuously for 14 days, and survival, behavioral changes, food intake, and body-weight changes were recorded.

At the end of the experiment, mice were euthanized after isoflurane anesthesia. Gross pathological examination was performed, major organs were weighed, and organ coefficients were calculated as follows:Organ coefficient (%) = organ weight / fasting body weight × 100.

#### 2.7.2 28-Day Repeated-Dose Oral Toxicity Test

The 28-day repeated-dose oral toxicity test was conducted according to the National Food Safety Standard: 28-Day Oral Toxicity Test (GB 15193.22-2014).

Forty SPF-grade SD rats (8 weeks old; half male and half female) were randomly assigned to 4 groups (n = 10 per group): control group, low-dose group (0.9 g/kg body weight), medium-dose group (1.8 g/kg body weight), and high-dose group (3.6 g/kg body weight). These doses were approximately equivalent to 5, 10, and 20 times the recommended human intake, respectively.

Animals were orally administered once daily for 28 consecutive days. General health status, food intake, and body weight were monitored during the experiment. At the end of the experiment, necropsy was performed; the heart, liver, spleen, kidney, and stomach were weighed; and organ coefficients were calculated.

Blood samples were collected for hematological analysis and serum biochemical testing, including aspartate aminotransferase (AST), alanine aminotransferase (ALT), albumin, total protein, creatinine, and urea nitrogen. Major organs were fixed, paraffin-embedded, sectioned, stained with hematoxylin and eosin (H&E), and examined histopathologically under a microscope. Representative H&E-stained sections from blank-control and high-dose animals were examined at ×20 and ×40 magnification. Male and female animals were evaluated separately. Histopathology was assessed qualitatively because no ordinal lesion-scoring data were available.

#### 2.7.3 In silico toxicological screening and network analysis

Putatively annotated MS/MS features were screened using predefined toxicology-related keyword categories, including mycotoxin-, pesticide/herbicide-, biogenic amine-, alkaloid/cyanogenic compound-, process-related-, and fermentation-related terms. Up to 12 matched features with the lowest Benjamini–Hochberg-adjusted P values in each category were visualized. The resulting toxicology watchlist, together with selected S7/DH signature features, comprised 68 candidates for prediction across 45 endpoints using ProTox 3.0 [38]. For visualization, model-reported probabilities were retained for endpoints predicted as active and set to zero for endpoints predicted as inactive.

Candidate names were matched against CTD chemical names, identifiers, curated synonyms, and aliases [39]. Of the 68 candidates, 56 were mapped to CTD, and only human (Homo sapiens) chemical–gene associations were retained. Genes were ranked first by the number of distinct mapped compounds and then by the total number of CTD interactions. Chemical–disease associations were grouped into nine predefined disease domains. Keyword matches, toxicity predictions, and CTD associations were interpreted as hypothesis-generating and not as confirmation of chemical identity, biological activity, or toxicological effects.

### 2.8 Establishment of the Gut Microbiota Dysbiosis Model and Intervention Evaluation

To evaluate the effects of WWPFS on gut microbiota dysbiosis, an Escherichia coli-induced mouse model of gut microbiota dysbiosis was established.

Male Kunming mice were randomly assigned to five groups: normal control group, model group, WWPFS high-dose group, WWPFS low-dose group, and positive control group. Except for the normal control group, mice were orally administered an E. coli suspension (2.45 × 10^8 CFU/mL; 15 mL/kg body weight) for 7 consecutive days to induce gut microbiota dysbiosis [40].

Intervention was initiated from the first day of model induction. The WWPFS high- and low-dose groups received WWPFS at 3.12 and 0.78 g/kg body weight, respectively. The positive control group received Bifidobacterium quadruple viable tablets (0.585 g/kg body weight), and the normal control and model groups received equal volumes of purified water.

After 14 consecutive days of intervention, animals were euthanized 24 h after the final administration. Colonic contents were aseptically collected and subjected to 16S rRNA sequencing to evaluate gut microbiota composition and the modulatory effects of WWPFS.

A Medium-dose group (n = 5) was also included but was excluded post hoc from the primary microbiome comparisons. Its sample-level sequencing and microbiome data are provided as an exploratory dataset in Supplementary Data 1..

### 2.9 Statistical Analysis

Statistical analyses were performed using SPSS 26.0 and R 4.5.2. Tests were two-sided unless otherwise stated. For Fig. 3a, total soluble solids, total carbon, total nitrogen, and the C/N ratio were compared between S1 and S3 using Welch’s t-tests based on three independent batches per stage. The four P values were adjusted using the Benjamini-Hochberg procedure.

**Figure 1.**
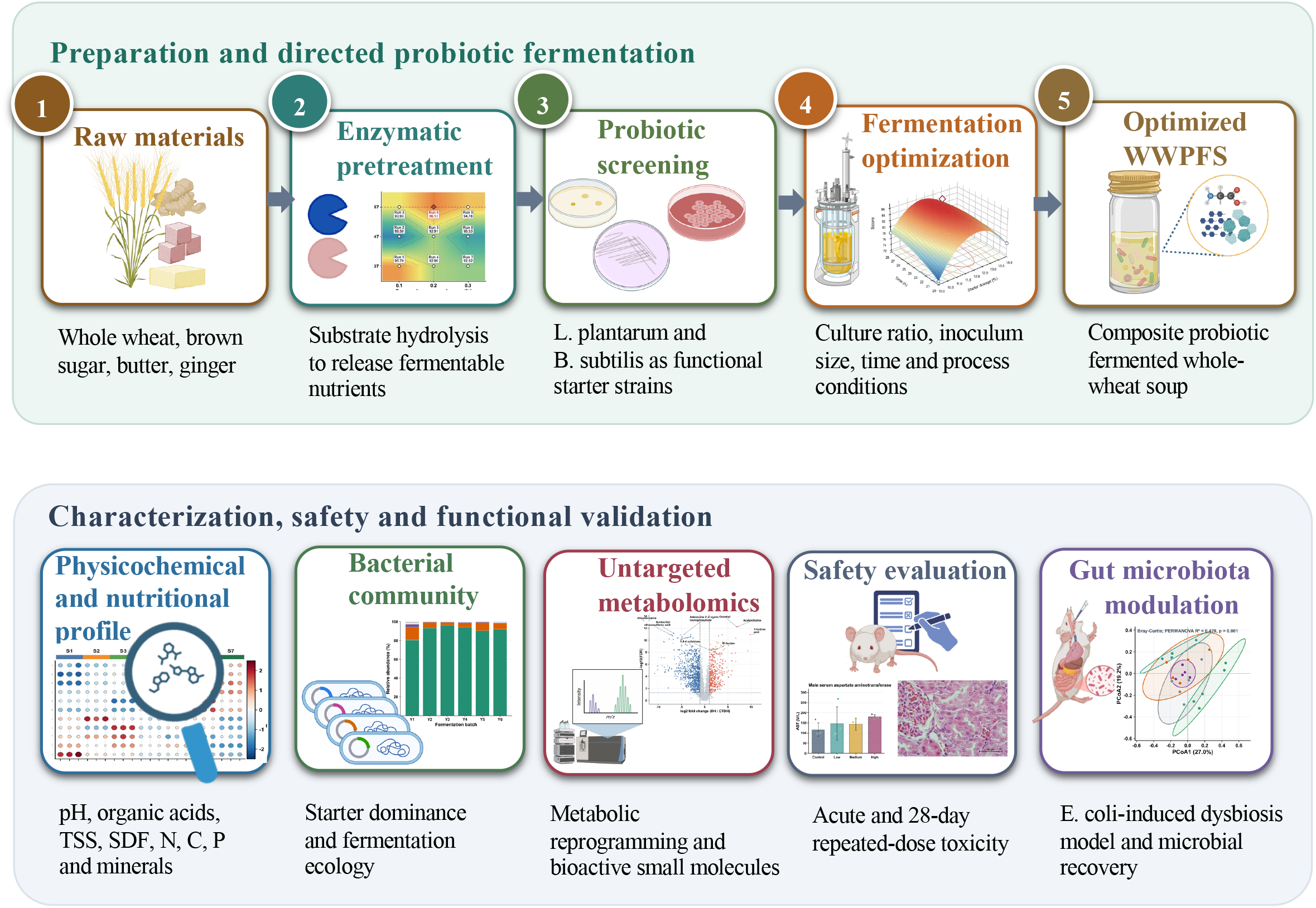
Workflow for the preparation, probiotic-directed fermentation, and biological evaluation of WWPFS. Schematic overview of WWPFS preparation and evaluation. Whole wheat-based medicinal dietary materials were processed by enzymatic pretreatment and probiotic-directed fermentation using Lactiplantibacillus plantarum and Bacillus subtilis as starter strains. The optimized WWPFS was characterized by physicochemical and nutritional analyses, bacterial community profiling, untargeted metabolomics, toxicological evaluation, and an E. coli-induced gut microbiota dysbiosis model.

**Figure 2.**
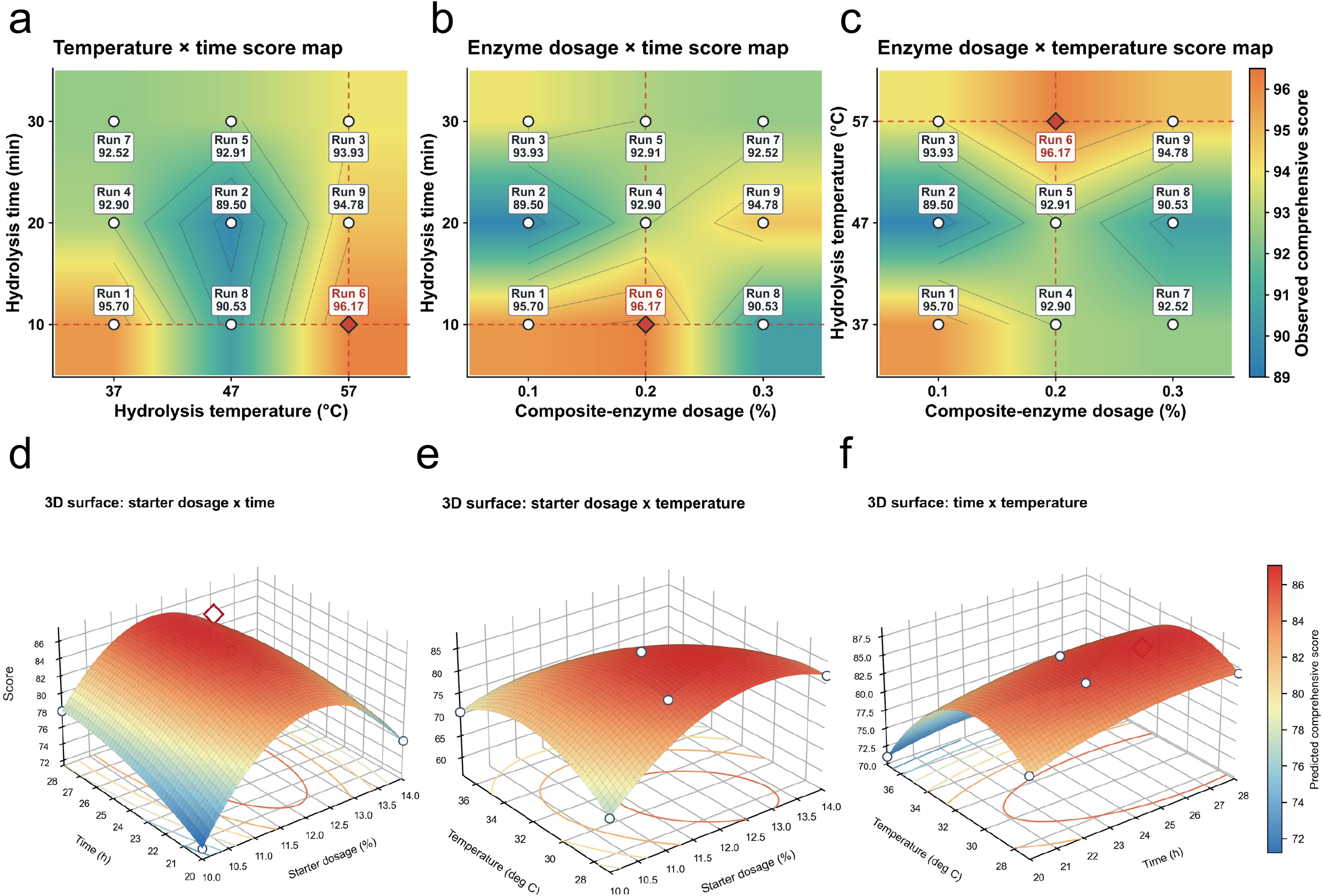
Optimization of enzymatic hydrolysis and probiotic fermentation conditions for WWPFS. Optimization of WWPFS processing conditions. (a–c) Score maps showing the composite hydrolysis scores across the tested factor combinations in the orthogonal design: (a) hydrolysis temperature × hydrolysis time; (b) total enzyme dosage × hydrolysis time; and (c) total enzyme dosage × hydrolysis temperature. (d–f) Three-dimensional response-surface plots showing the combined effects of fermentation parameters on the predicted composite score. d) Starter dosage (hereafter using inoculum size) × fermentation time; (e) inoculum size × fermentation temperature; and (f) fermentation time × fermentation temperature. The apex of each 3D surface (indicated by a red diamond) represents the predicted maximum response, indicating the optimal combination of variables.

**Figure 3.**
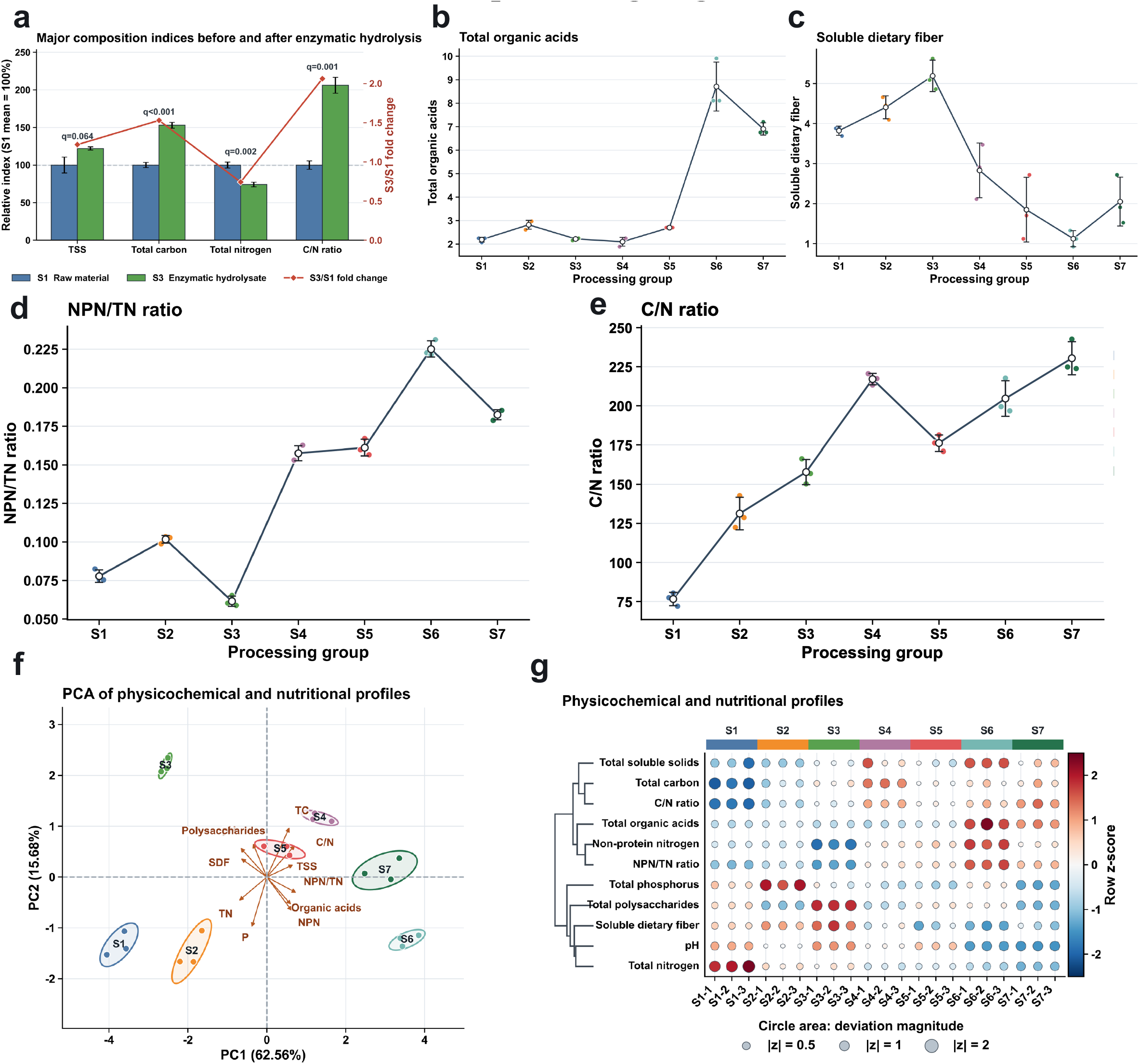
Physicochemical and nutritional remodeling of WWPFS across processing stages. Effects of enzymatic pretreatment and probiotic fermentation on the physicochemical and nutritional characteristics of WWPFS. (a) Changes in major composition indices (TSS, Total carbon, Total nitrogen, and C/N ratio) before and after composite enzymatic hydrolysis (S1 raw material vs. S3 hydrolysate). Bars represent the relative index (%), and the red line denotes the S3/S1 fold change (q-values represent FDR-adjusted statistical significance). (b) Total organic acid accumulation across processing groups (S1– S7). (c) Differences in soluble dietary fiber (SDF) content across processing groups. (d) Non-protein nitrogen (NPN/TN ratio) content across processing groups, highlighting the small-molecule nitrogen transformation during fermentation. **(e)** Shift in the carbon-to-nitrogen (C/N ratio) during different fermentation schemes. **(f)** Principal component analysis (PCA) score and loading biplot based on combined physicochemical and nutritional indicators, illustrating distinct group separations (PC1 = 62.56%, PC2 = 15.68%). **(g)** Clustered heatmap displaying normalized row Z-scores of 11 primary physicochemical parameters across all replicates . Circle sizes and colors indicate deviation magnitude and expression direction, respectively. (Processing group definitions: S1, Raw material; S2, Traditional fermentation; S3, Enzymatic hydrolysate; S4, Sterilized control; S5, B. subtilis fermentation; S6, L. plantarum fermentation; S7, Mixed-probiotic fermentation).

Microbiome alpha-diversity indices, including observed OTUs, Shannon diversity, Simpson diversity, Chao1 richness, and Good’s coverage, were recalculated from the provider-supplied annotated OTU count table. Overall differences were assessed using Kruskal-Wallis tests, followed by pairwise Wilcoxon rank-sum tests with Benjamini-Hochberg correction within each index [41]. Alpha-rarefaction curves were reconstructed using hypergeometric expected richness at 14 depths from 1,000 to 50,000 retained sequence tags per sample.

Bray-Curtis dissimilarities [42] were calculated from OTU-level relative abundances and visualized by PCoA using classical multidimensional scaling. Global differences were assessed using PERMANOVA with 999 permutations [43]. PERMDISP was performed from distances to group spatial medians using vegan::betadisper and 999 permutations. Four Model-centred pairwise comparisons against the Blank, Low-dose, High-dose, and Positive-control groups used vegan::adonis2 with 999 permutations. The four P values were adjusted using the Benjamini-Hochberg procedure.

For genus-level analysis, OTU abundances were aggregated using the provider-assigned genus annotations. Model-versus-Blank differences were tested using Wilcoxon rank-sum tests. Log2 fold changes were calculated as log2[(Model mean + 0.001)/(Blank mean + 0.001)], where abundance was expressed in percentage points and 0.001 was used as a pseudocount. P values were adjusted across all tested genera using the Benjamini-Hochberg procedure. Genera with nominal P < 0.05 were displayed for exploratory visualization.

For the exploratory rescue analysis, the treatment-specific rescue score was calculated as:

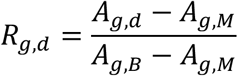

where g denotes the genus and d denotes the Low-dose, High-dose, or Positive-control group. Scores were treated as missing when the denominator magnitude was below 1 × 10−9 and were otherwise truncated to −1.5 to 1.5. The primary rescue score was the mean of the Low- and High-dose scores, excluding the Positive-control score, and was displayed on a scale from 0 to 1.5.

For untargeted LC-MS/MS data, non-positive intensities were replaced with one-half of the smallest positive value per feature before log2 transformation. S7 and S2 were compared using Welch’s t-tests, with Wilcoxon rank-sum tests as sensitivity analyses. Benjamini-Hochberg correction was applied across features, and differential features were defined as q < 0.05 and |log2 fold change| ≥ 1. PCA used feature-wise standardization, and Spearman correlations were calculated in R.

Putatively annotated differential features were mapped to KEGG pathways and evaluated using an upper-tail hypergeometric test against all retained KEGG-annotated features. The enrichment ratio was defined as the pathway proportion among differential features divided by the corresponding background proportion. Pathway P values were Benjamini-Hochberg-adjusted; Fig. 5d reports nominal P values and was interpreted as exploratory.

**Figure 4.**
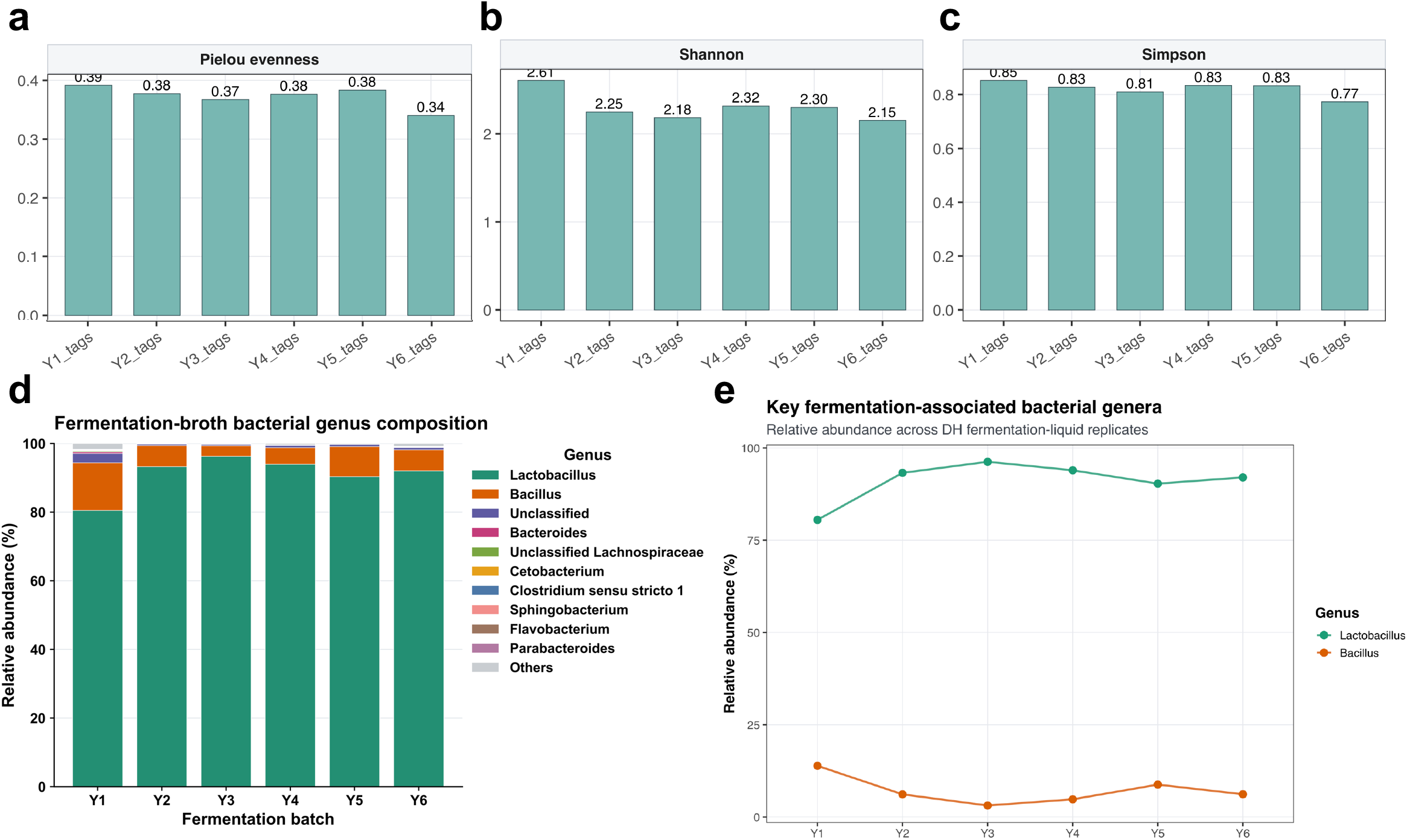
Bacterial diversity and genus-level community structure across WWPFS fermentation batches. Bacterial community profiling of fermentation-liquid replicates. (a) Pielou evenness. (b) Shannon diversity. (c) Simpson diversity. (d) Genus-level microbial composition across fermentation batches. (e) Relative abundances of Lactobacillus and Bacillus.

**Figure 5.**
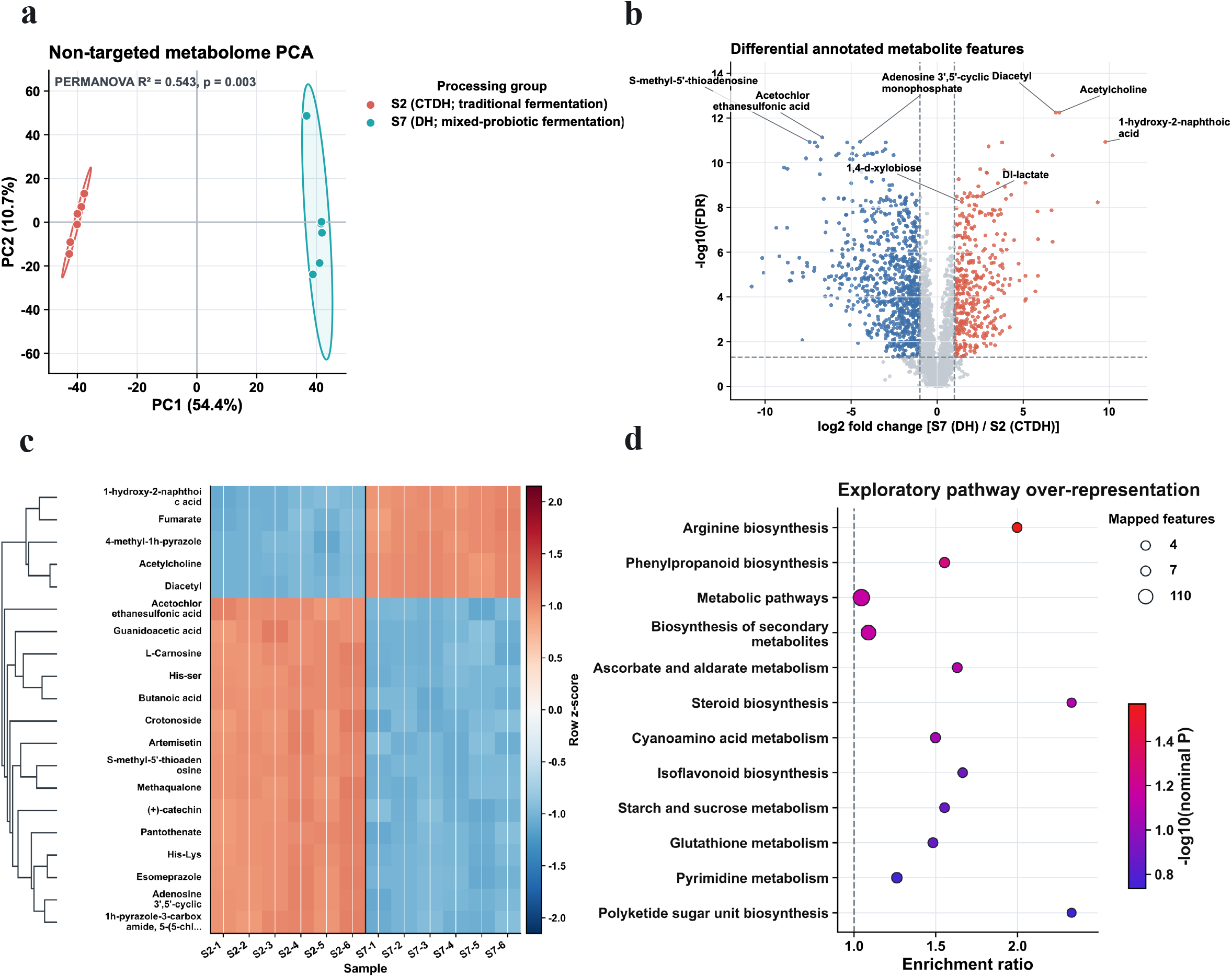
Probiotic-directed fermentation reshapes the WWPFS metabolite profile compared with traditional processing. Untargeted metabolomic comparison between traditional fermentation (S2) and probiotic-directed fermentation (S7). (a) PCA score plot showing separation between S2 and S7 samples with tight clustering of biological replicates. (b) Volcano plot showing differential annotated metabolite features between S7 and S2 , including representative S7-enriched features such as lactate, phenyllactic acid- related metabolites, diacetyl-associated features, and xylobiose-related. (c) Hierarchical clustering heatmap confirming process-dependent metabolite-pattern differences. (d) Pathway enrichment analysis of differential features, highlighting metabolic shifts related to carbohydrate conversion, organic acid formation, amino acid-associated metabolism, and flavor-related small molecules.

Acute organ coefficients were analyzed separately by sex using Welch’s t-tests, with Benjamini-Hochberg correction across five organs within each sex. Body weight was summarized descriptively by group, sex, and study week as mean ± SD.

Repeated-dose hematological endpoints were analyzed separately by sex using Kruskal-Wallis tests, with correction across eight endpoints within each sex. Exploratory dose-versus-control comparisons used Mann-Whitney U tests with Benjamini-Hochberg correction across all 48 sex-specific comparisons. Hematological values were displayed as fold changes relative to sex- matched controls. Histopathological findings were evaluated qualitatively without inferential testing.

Adjusted q < 0.05 was considered statistically significant. Nominal P < 0.05 was used only for explicitly exploratory analyses. Figures were generated using GraphPad Prism 10.0 and R 4.5.2.

## 3. Results

### 3.1 Optimization of WWPFS Fermentation Conditions

Phenolic acids and flavonoids are important bioactive markers in the whole-wheat and dried- ginger matrix. Their release during enzymatic hydrolysis and subsequent microbial transformation can reflect, respectively, substrate accessibility and fermentation efficiency [20,44–46]. Therefore, total polyphenol content, total flavonoid content, and viable cell count were selected as integrated endpoints to optimize the two-stage process of enzymatic hydrolysis followed by probiotic-directed fermentation.

Under fixed pH and cellulase-to-pectinase ratio conditions, single-factor experiments and orthogonal design analysis showed that total flavonoid and total polyphenol release were highest when the hydrolysis temperature was 57 °C, hydrolysis time was 10 min, and total enzyme dosage was 0.2%. In validation experiments, the measured values did not differ significantly from the predicted values (P > 0.05), supporting the reproducibility of the optimized enzymatic hydrolysis process (Fig. 2a-c; Supplementary Fig. S1c-f; Supplementary Tables S2 and S3).

Response surface methodology was subsequently used to optimize the fermentation stage. The quadratic response-surface model was significant (F = 19.86, P = 0.0003), with an R² of 0.9623 and an adjusted R² of 0.9139. The lack-of-fit test was not significant (P = 0.9134). Fermentation temperature (C), the interaction between inoculum size and fermentation temperature (AC), and the quadratic terms A² and C² were significant, whereas the AB and BC interaction terms were not significant. Among the experimental runs, the highest observed composite score was 90.98 at an inoculum size of 12%, fermentation time of 24 h, and fermentation temperature of 32 °C. Based on the fitted model, the predicted optimum was an inoculum size of 12.29%, fermentation time of 25.83 h, and fermentation temperature of 29.70 °C (Fig. 2d–f; Supplementary Fig. S1n–p; Supplementary Tables S5 and S6).

### 3.2 Probiotic-Directed Fermentation Reshaped the Physicochemical and Nutritional Features of WWPFS

To clarify how enzymatic pretreatment and probiotic-directed fermentation jointly shaped WWPFS quality, physicochemical and nutritional indices were compared across seven treatment systems. The analysis was organized around a sequential logic: matrix deconstruction, fermentable carbon release, acid and nitrogen conversion, and integrated nutritional remodeling.

Enzymatic pretreatment altered the physicochemical composition of the raw-material matrix. Compared with the raw-material group (S1), the enzymatic hydrolysate group (S3) showed higher TSS, total carbon, and polysaccharide contents, together with lower total nitrogen. These changes indicate that cellulase-pectinase treatment loosened cereal cell-wall and polysaccharide networks and shifted the hydrolysate toward a carbon-enriched substrate pool, thereby increasing the availability of soluble carbohydrate resources for downstream microbial metabolism (Fig. 3 a).

Viable fermentation then redirected this carbon-enriched hydrolysate toward secondary microbial conversion. Compared with S3 and the heat-inactivated control (S4), S5-S7 showed lower pH and higher organic acid levels, indicating that active microbial metabolism was a major contributor to the observed acidification. The probiotic consortium group (S7) combined evident acidification with increased NPN/TN, suggesting that carbon utilization and the formation or release of non-protein nitrogen components occurred in parallel. (Fig. 3b-d).

Elemental analysis further showed that fermentation reshaped the carbon-mineral balance of WWPFS. S7 displayed a high C/N ratio and relatively coordinated Mg, Mn, and sulfur-related profiles, suggesting that consortium fermentation did not simply intensify a single pathway, but produced a balanced nutritional transformation involving carbon redistribution, nitrogen fractionation, and elemental composition (Fig. 3e; Supplementary Fig. S2 c).

PCA integrated these variables and separated the seven treatments according to their overall quality profiles. S6 was mainly associated with stronger acidification and NPN formation, whereas S7 occupied a distinct region linked to C/N balance and consortium-driven transformation. Thus, the advantage of the probiotic consortium was not the maximum value of every index, but an integrated profile that connected substrate deconstruction, carbon flow, acidification, nitrogen redistribution, and elemental composition (Fig. 3 f. g).

### 3.3 Optimized Fermentation Established a Reproducible Probiotic-Dominated Community across Independent Batches

To clarify the microbial basis underlying the physicochemical changes described above, high-throughput sequencing was used to characterize the bacterial community structure of WWPFS fermentation products. Across the six independent fermentation batches, Pielou evenness ranged from 0.340 to 0.392, Shannon diversity from 2.151 to 2.606, and Simpson diversity from 0.773 to 0.853 (Fig. 4a–c). Lactobacillus was the predominant genus, ranging from 80.5% to 96.2%, whereas Bacillus ranged from 3.1% to 13.9% across batches (Fig. 4d, e).

### 3.4 Probiotic-Directed Fermentation Drove Metabolic Reprogramming of WWPFS

Based on the stable dominance of the screened probiotic strains and the physicochemical evidence of carbon and nitrogen redistribution, untargeted LC–MS/MS metabolomics was used to determine whether these bulk compositional changes were accompanied by small-molecule metabolic remodeling in WWPFS.

PCA showed clear separation between the traditional fermentation group (S2) and the probiotic fermentation group (S7), whereas biological replicates within each group clustered tightly (Fig. 5a; Supplementary Fig. S3). Volcano plot analysis further showed that probiotic- directed fermentation generated a distinct set of differential metabolite features, with representative S7-enriched features including acetylcholine, diacetyl, DL-lactate, fumarate, and 1,4-D-xylobiose (Fig. 5b). Hierarchical clustering confirmed that the two fermentation strategies generated distinct metabolite profiles (Fig. 5c), supporting a process-dependent rather than random compositional shift.

Differential feature analysis showed that features putatively annotated as acetylcholine, DL-lactate, 1,4-D-xylobiose, diacetyl, and fumarate showed higher relative signal intensities in the S7 group. In contrast, features putatively annotated as adenosine, deoxyadenosine, S-methyl-5′- thioadenosine, and several precursor-like nitrogenous features were higher in the S2 group (Fig. 5b-c; Supplementary Fig. S3). Among the increased S7 features, acetylcholine-related compounds, lactic acid, and xylobiose showed the most significant increases (FDR < 0.001).

Pathway over-representation analysis mapped the putatively annotated differential features mainly to starch and sucrose metabolism, arginine biosynthesis, glutathione metabolism, phenylpropanoid biosynthesis, and other secondary-metabolite-related pathways (Fig. 5d). However, no pathway remained significant after Benjamini-Hochberg correction (minimum q = 0.962). These nominal pathway-level patterns were therefore considered exploratory and were not interpreted as evidence of confirmed pathway activation. At the individual-feature level, the S7 profile was nevertheless characterized by higher relative signals for several putatively annotated organic-acid-, oligosaccharide-, and aroma-related features.

### 3.5 Safety Evaluation Revealed No Treatment-Related Adverse Effects under the Tested Conditions

To evaluate the safety of WWPFS after probiotic-directed fermentation, acute oral toxicity and 28-day repeated-dose toxicity tests were performed.

In the acute oral toxicity test, no mortality, behavioral abnormalities, or obvious toxic manifestations were observed after high-dose WWPFS administration. During the observation period, food intake, activity, and body-weight gain remained normal. Gross examination showed no obvious abnormalities in major organs, and organ coefficients were within the normal range (Fig. 6a, b).

**Figure 6.**
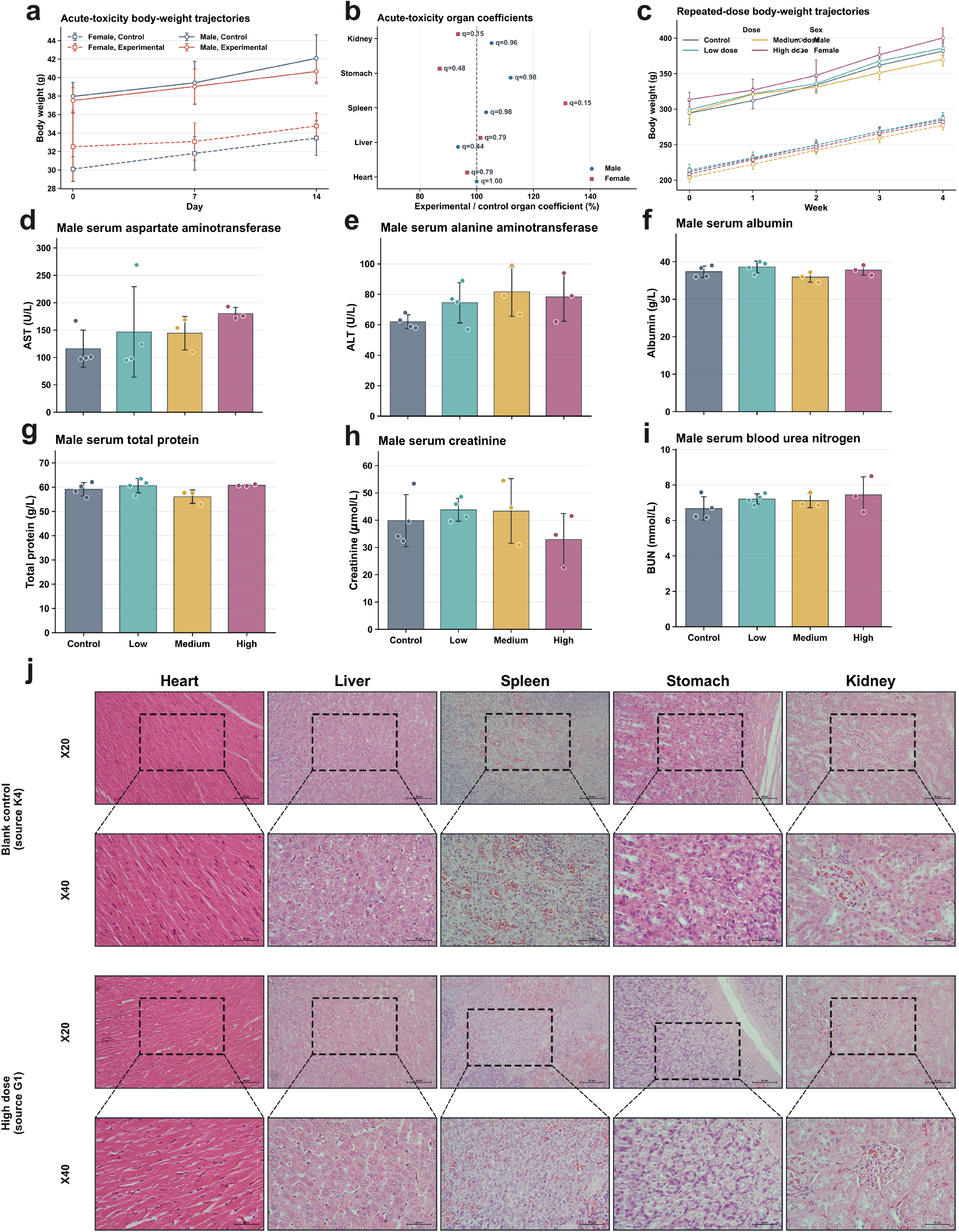
In vivo toxicological evaluation of WWPFS in acute and 28- day repeated-dose oral toxicity studies. Safety assessment of WWPFS in mice and rats. (a) Body-weight trajectories during the acute toxicity study. (b) Relative organ coefficients after acute administration. (c) Body-weight trajectories of male and female rats during the 28-day repeated-dose study. (d–i) Serum biochemical parameters in male rats: (d) AST, (e) ALT, (f) albumin, (g) total protein, (h) creatinine, and (i) blood urea nitrogen. (j) Representative H&E-stained sections of the heart, liver, spleen, stomach, and kidney from control and high-dose rats.

Similarly, in the 28-day repeated-dose toxicity test, body weight increased normally in all administration groups, with no significant difference compared with the control group (P > 0.05). Organ morphology, organ coefficients, hematological parameters, serum biochemical indices, and histopathological examination showed no treatment-related abnormalities (Fig. 6c–j; Supplementary Fig. S4a, b). These results indicate that WWPFS did not show observable acute or repeated-dose toxicity within the tested dose range.

Repeated-dose hematological profiles were broadly comparable with sex-matched controls, with no significant overall differences after correction across endpoints (all q > 0.05), and qualitative histological examination did not reveal overt high-dose-associated lesions (Supplementary Fig. S4a, b). To complement these findings, differential features from untargeted metabolomics were subjected to keyword-based toxicological screening, ProTox 3.0 prediction, and CTD network analysis. The S7/DH profile was dominated by putatively annotated fermentation-related features and showed generally lower predicted toxicological activity than the toxicology-watchlist profile, whereas several features enriched in S2/CTDH matched categories requiring targeted confirmation (Supplementary Fig. S4c–f). CTD analysis further prioritized candidate hub genes and disease-associated domains. Because metabolite annotations were putative and the network analyses were computational, these findings were treated as hypothesis-generating rather than independent evidence of chemical identity, in vivo toxicity, or product safety.

### 3.6 WWPFS Was Associated with Gut Microbiota Shifts in an Escherichia coli-Induced Dysbiosis Model

An Escherichia coli-induced dysbiosis model was used to evaluate the effects of WWPFS intervention on mouse gut microbial diversity and community composition.

Overall differences in alpha diversity were observed among groups (Kruskal-Wallis: observed OTUs, P = 0.008; Shannon diversity, P = 0.023; Simpson diversity, P = 0.031; and Chao1 richness, P = 0.007). The High-dose group showed numerically higher alpha-diversity values, but pairwise comparisons did not remain significant after multiple-testing correction (Fig. 7a).

**Figure 7.**
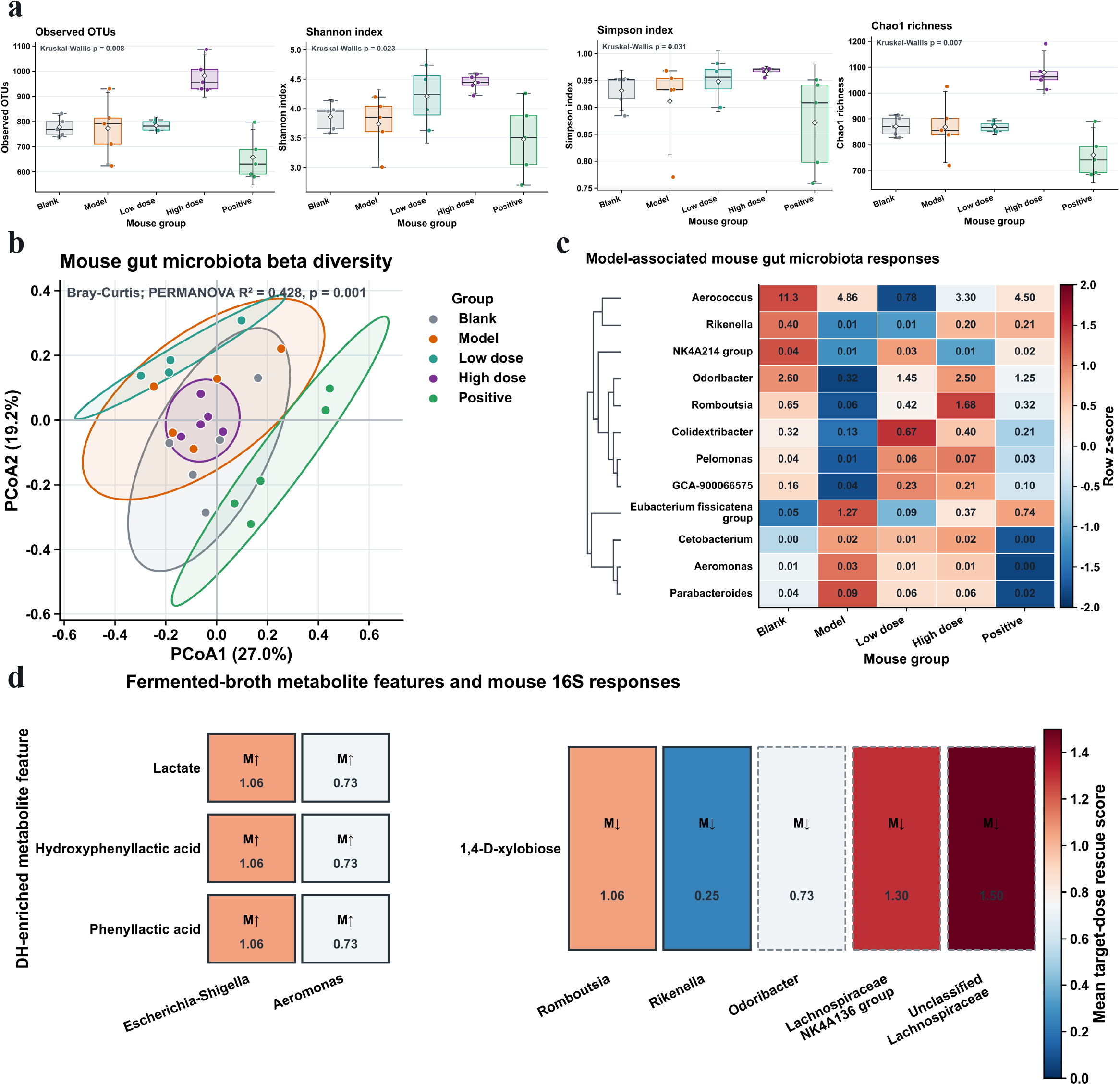
Gut microbiota modulation and metabolite-microbiome response alignment in mice receiving WWPFS. Gut microbiota modulation in mice receiving WWPFS. (a) Alpha-diversity metrics, including observed OTUs, Shannon index, Simpson index, and Chao1 richness. (b) Principal coordinate analysis based on Bray-Curtis distances showing group-level microbial community separation. (c) Heatmap of representative model-responsive bacterial genera across the five experimental groups. (d) Cross-system response-alignment analysis linking fermentation-derived metabolite features with directional rescue of model-responsive gut bacterial taxa. The scores in squares are taxon rescue score. M↑, increased in the model group relative to blank; M↓, decreased in the model group relative to blank. Solid outline: observed microbial recovery pattern Dashed outline: literature-based candidate mechanism

Bray-Curtis PCoA showed a significant global group effect (PERMANOVA, R² = 0.428, P = 0.001), although partial overlap remained among the Blank, Model, and High-dose groups (Fig. 7b). Multivariate dispersion did not differ significantly among groups (PERMDISP, F = 1.155, P = 0.351), providing no evidence that the global PERMANOVA result was primarily attributable to unequal within-group dispersion. In the four Model-centred pairwise comparisons, the Low-dose, High-dose, and Positive-control groups differed significantly from the Model group after Benjamini-Hochberg correction (Model versus Low: R² = 0.276, P = 0.007, q = 0.0147; Model versus High: R² = 0.239, P = 0.008, q = 0.0147; and Model versus Positive: R² = 0.339, P = 0.011, q = 0.0147). The Model-versus-Blank comparison did not reach significance after adjustment (R² = 0.203, P = 0.053, q = 0.053).

Several genera showed nominal differences between the Model and Blank groups; however, none remained significant after Benjamini-Hochberg correction (all q ≥ 0.350; Supplementary Fig. S5b). Exploratory genus-level comparisons showed higher mean abundances of Aeromonas, Cetobacterium, and Parabacteroides in the Model group, whereas Rikenella and Odoribacter showed lower mean abundances. WWPFS-treated groups showed directional shifts in these genera toward the corresponding Blank-group means, but these genus-level patterns were not statistically significant after multiple-testing correction (Fig. 7c; Supplementary Fig. S5b). The associated taxon rescue scores were therefore considered exploratory.

An exploratory cross-system response-alignment analysis compared the directions of S7- versus-S2 metabolomic changes with the genus-level shifts observed after WWPFS intervention. Lactate-, phenyllactic-acid-, and hydroxyphenyllactic-acid-related features showed directional correspondence with lower mean abundances of Escherichia-Shigella and Aeromonas, whereas 1,4-D-xylobiose showed directional correspondence with shifts in Romboutsia, Rikenella, Odoribacter, and Lachnospiraceae-related taxa (Fig. 7d). Because the metabolomic and mouse microbiome datasets were generated from separate experimental systems, this analysis did not test direct metabolite-microbiota associations.

Together, WWPFS intervention was associated with changes in overall gut microbial community structure in the Escherichia coli-induced model. The genus-level rescue patterns and cross-system metabolite-microbiota alignment remain exploratory and require validation using targeted metabolite quantification, strain-resolved microbiome analysis, and direct supplementation experiments.

## 4. Discussion

This study shows that the main value of probiotic-directed fermentation in WWPFS is controlled metabolic reprogramming of the whole-wheat matrix. Unlike spontaneous fermentation, the defined Lactobacillus-Bacillus starter improved process controllability, microbial dominance, and substrate conversion. More importantly, the study links physicochemical remodeling with untargeted metabolomic changes, providing a mechanistic bridge from matrix deconstruction to potential gut microbiota modulation.

Although *L. plantarum* and *B. subtilis* were inoculated at a 1:1 ratio, the Lactobacillus-dominated structure after fermentation is consistent with the combined selection imposed by substrate availability, acidification, and oxygen availability. Whole-wheat hydrolysate and brown sugar supplied readily fermentable carbohydrates, favoring rapid growth and organic acid production by *L. plantarum*. The vessels were sealed with a microporous plant tissue culture film and aluminum foil and incubated without forced aeration, providing limited gas exchange rather than active aeration. Together with the observed pH decrease and organic acid accumulation, this condition would favor acid-tolerant lactic acid bacteria while restricting sustained proliferation of vegetative *B. subtilis* cells. This bacterium may therefore contribute mainly during the early stage by secreting extracellular enzymes that open the cereal matrix and provide accessible substrates for *L. plantarum*. Thus, the final microbial structure reflects substrate preference, acid tolerance, oxygen adaptation, and stage-dependent ecological selection rather than the initial inoculation ratio alone [48–50].

The consortium advantage is best interpreted as functional division rather than universal maximization of all indices. *B. subtilis* can contribute extracellular enzymes that open cell-wall and protein-polysaccharide structures, while *L. plantarum* drives acidification and stabilizes the fermentation ecology. This division explains why S7 showed a balanced pattern of carbon availability, nitrogen redistribution, organic acid formation, and elemental composition instead of simply reproducing the strongest single-strain response. In this sense, S7 represents a coordinated biotransformation mode rather than an extreme acidification mode [47–50].

Untargeted metabolomics further supports this function-oriented shift and provides the key link between bulk physicochemical indices and molecular function. Enrichment of lactic acid, xylobiose, diacetyl, phenyllactic acid, and hydroxyphenyllactic acid links carbohydrate utilization, hemicellulose-derived oligosaccharide release, flavor formation, and antimicrobial potential. Therefore, the innovation of WWPFS fermentation is not only the selection of probiotic strains, but also the establishment of a sequential enzymatic degradation-probiotic biotransformation cascade that converts a cereal matrix into a small-molecule-enriched fermented system [51–54].

Safety evaluation supports the edible application potential of WWPFS within the tested dose range. The Lactobacillus-dominated community, acidified environment, and antimicrobial organic-acid derivatives may also reduce contamination risk during controlled fermentation. Nevertheless, future food-safety assessment should include targeted monitoring of biogenic amines, mycotoxins, viable spore persistence, antimicrobial-resistance genes, and long-term intake safety.

Functionally, WWPFS alleviated E. coli-induced gut microbiota dysbiosis through a likely multi-component mode of action. Lactic acid may act as a cross-feeding substrate for short-chain fatty acid formation, xylo-oligosaccharide-like compounds may support beneficial taxa, and phenyllactic-acid-related metabolites may suppress opportunistic bacteria. These linked effects are consistent with the observed partial restoration of Rikenella, Odoribacter, Romboutsia, and Lachnospiraceae-related taxa [52–54]. Because fermentation metabolomics and mouse gut microbiota profiles were obtained from separate experimental systems, these cross-system patterns should be interpreted as mechanistic hypotheses rather than direct metabolite– microbiota associations.

The exploratory Medium-dose group showed substantial inter-individual variability, with Escherichia-Shigella relative abundance ranging from 1.4% to 80.3% (Supplementary Data 1) and did not exhibit a consistent treatment-associated microbiota response. Because 16S rRNA gene sequencing does not resolve strain-level engraftment, this pattern cannot be interpreted as confirmed colonization failure. The post hoc exclusion of this group from the primary comparisons should therefore be considered a limitation of the microbiome analysis.

Overall, probiotic-directed fermentation transformed WWPFS through a sequence of matrix opening, carbon mobilization, acidification, and metabolite enrichment, followed by evaluation of its safety and gut microbiota-modulating effects. The Lactobacillus-Bacillus consortium therefore provides a rational strategy for converting a traditional whole-wheat preparation into a more controllable fermented food system. Future studies should verify strain-level dynamics, enzyme activities, targeted metabolites, mineral bioaccessibility, and causal gut-microbiota mechanisms.

## 5. Conclusion

This study established a controllable probiotic-directed fermentation process for a traditional Mongolian whole-wheat dietary preparation. Enzymatic pretreatment improved substrate accessibility, and the selected Lactobacillus-Bacillus consortium formed a stable fermentation system accompanied by physicochemical transformation, metabolic reprogramming, and enrichment of fermentation-derived small molecules. No treatment-related adverse effects were observed within the tested dose range and duration, and WWPFS intervention was associated with partial alleviation of E. coli-induced gut microbiota dysbiosis in mice. These results support probiotic-directed fermentation as a strategy for standardizing and modernizing traditional Mongolian medicinal fermented foods and for developing gut microbiota-modulating functional foods. Further targeted metabolomics and mechanistic studies are needed to validate the causal roles of key metabolites and microbial taxa.

### Use of generative artificial intelligence

OpenAI Codex was used to assist with English-language editing and development of data-analysis scripts. All analytical decisions, code, statistical outputs, and manuscript text were reviewed and verified by the authors, who take full responsibility for the reported work. OCEAN (https://github.com/nslbotnslbot/ocean-skill), an AI-assisted evidence-navigation workflow, was used to support literature retrieval, claim–evidence organization and citation checking. The authors reviewed the source literature, executed and validated the code and analyses, verified all citations and text, made all scientific and clinical decisions, and take full responsibility for the final manuscript.

## Supporting information

Medium_Dose_Microbiome

## Acknowledgements

This work was supported by the 2025 Key Discipline Construction Project of Traditional Chinese Medicine (Mongolian Medicine) of Inner Mongolia Autonomous Region, Mongolian Medicine Hematology, and the Graduate Research Innovation Project of Inner Mongolia Minzu University, “Changes in Functional Components and Metabolic Characteristics of Whole-Wheat Fermented Soup during Probiotic Fermentation.” The funders had no role in study design, data collection, data analysis, interpretation, or preparation of the manuscript.

## Data Availability

Raw 16S rRNA gene sequencing reads generated from the WWPFS fermentation-batch and mouse gut microbiota experiments are being deposited in the NCBI Sequence Read Archive. Raw LC–MS/MS data and the corresponding processed feature-intensity matrices are being deposited in MetaboLights. Repository accession numbers will be added in the next version of this preprint. Processed OTU tables, sample metadata, metabolomic feature tables, and source data underlying the figures are provided in the Supplementary Data files.

## CRediT Authorship Contribution Statement

Eerdunduleng: Conceptualization, Methodology, Investigation, Formal analysis, Writing - review & editing, Project administration, Funding acquisition.

Qingchun Ling: Investigation, Data curation, Formal analysis, Writing - original draft.

Jinhua Bao: Investigation, Formal analysis, Writing - original draft, Funding acquisition.

Surigaga: Investigation, Physicochemical analysis, Metabolomic analysis, Writing - original draft.

Tegexi: Investigation, Toxicological evaluation, Functional evaluation, Writing - original draft. Nandien: Investigation, WWPFS preparation, Writing - original draft.

Shandan: Data curation, Formal analysis, Writing - original draft, Writing - review & editing. Lisha A: Investigation, Data curation, Formal analysis, Writing - review & editing.

Cuiqin Gong: Conceptualization, Methodology, Writing - original draft, Funding acquisition. Burenbatu: Conceptualization, Methodology, Writing - review & editing.

Shengliang Ni: Formal analysis, Methodology, Data curation, Visualization, Writing – review & editing, Project administration.

Wenping: Conceptualization, Methodology, Investigation, Writing - review & editing, Project administration, Funding acquisition.

---

**Fig. S1.**
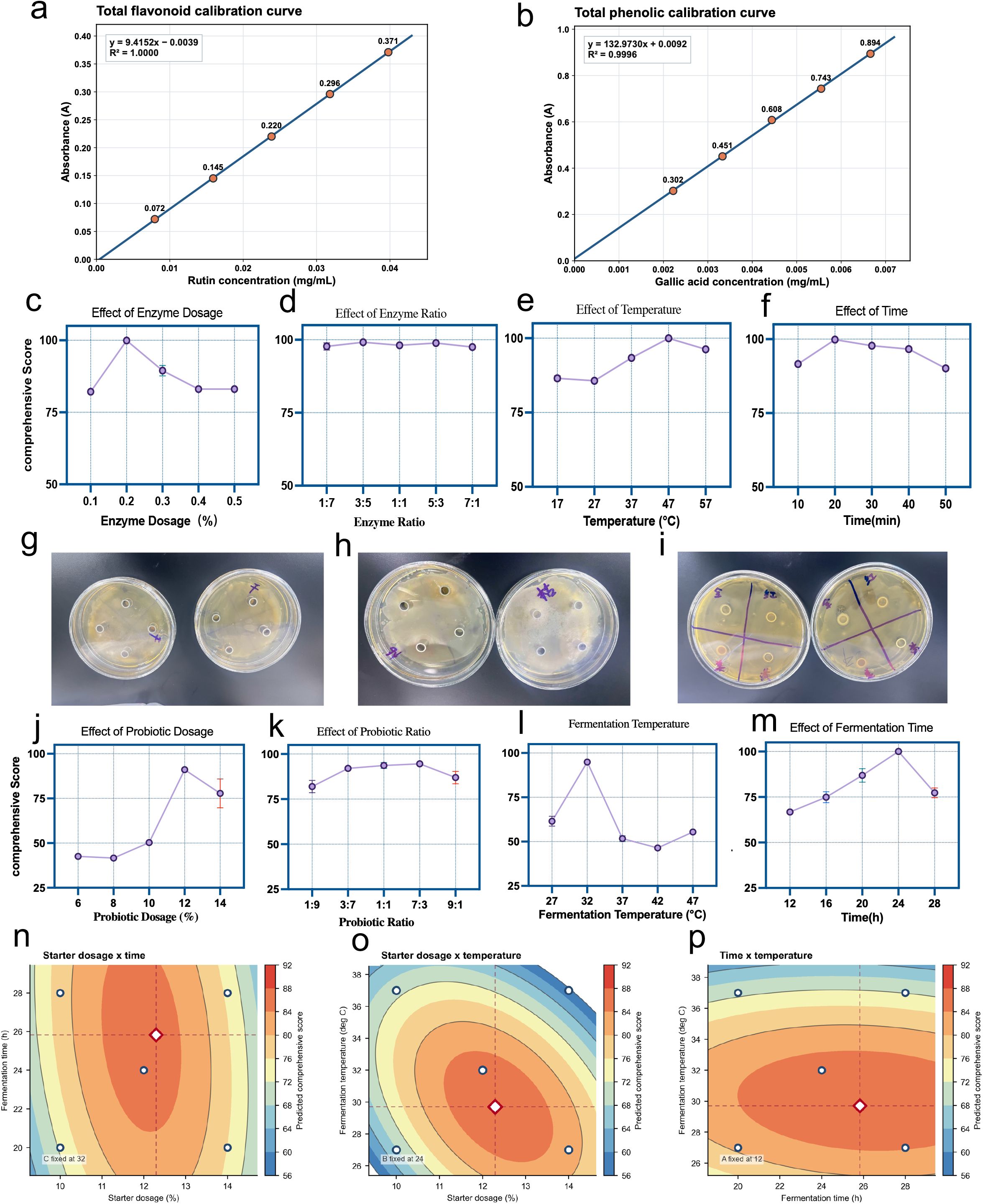
Method validation, single-factor screening, antibacterial assays, and response surface modeling for process optimization. (a, b) Standard calibration curves for quantification of functional components: (a) Total flavonoids (Rutin standard, y = 9.4152x − 0.0039, R^2^ = 1.0000) and (b) Total phenolic (Gallic acid, y = 132.973x + 0.0092, R^2^ = 0.9996). (c) Effect of total enzyme dosage on the composite hydrolysis score. (d) Effect of the cellulase-to-pectinase ratio. (e) Effect of hydrolysis temperature on the fermentation composite score. (f) Effect of hydrolysis time. (g-i) Representative agar diffusion assay plates showing in vitro antibacterial inhibition zones produced by Lactiplantibacillus plantarum (g), Bacillus subtilis (h), and Lacticaseibacillus casei (i) against target opportunistic pathogens. (j) Effect of probiotic dosage (equal to inoculum size). (k) Effect of probiotic ratio (L. plantarum to B. subtilis ratio). (l) Effect of fermentation temperature. (m) Effect of fermentation time. (n–p) Two-dimensional RSM contour plots depicting pairwise interactive effects on the predicted comprehensive score:(n) Starter dosage vs. Fermentation time; (o) Starter dosage vs. Fermentation temperature ; (p) Fermentation time vs. Fermentation temperature. Central diamonds indicate predicted global optimal domain.

**Supplementary Fig. S2.**
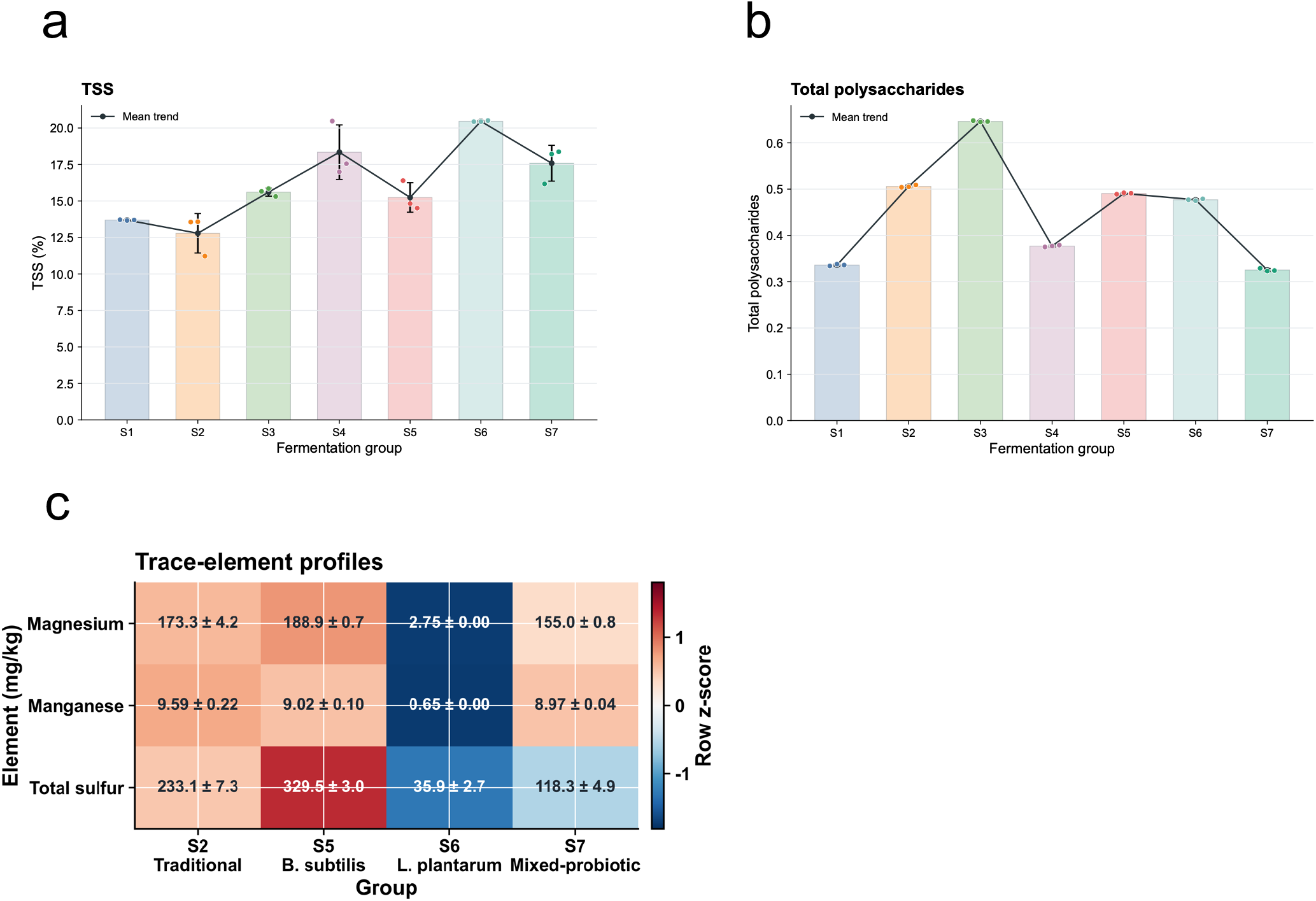
Supplementary physicochemical characteristics of WWPFS. (a) Total soluble solids (TSS, %) content across different processing groups. (b) Total polysaccharides content across different processing groups. (c) Heatmap comparing key trace elements (Magnesium, Manganese) and Total sulfur content (mg/kg) among representative fermentation groups (S2, S5, S6, and S7).

**Supplementary Fig. S3.**
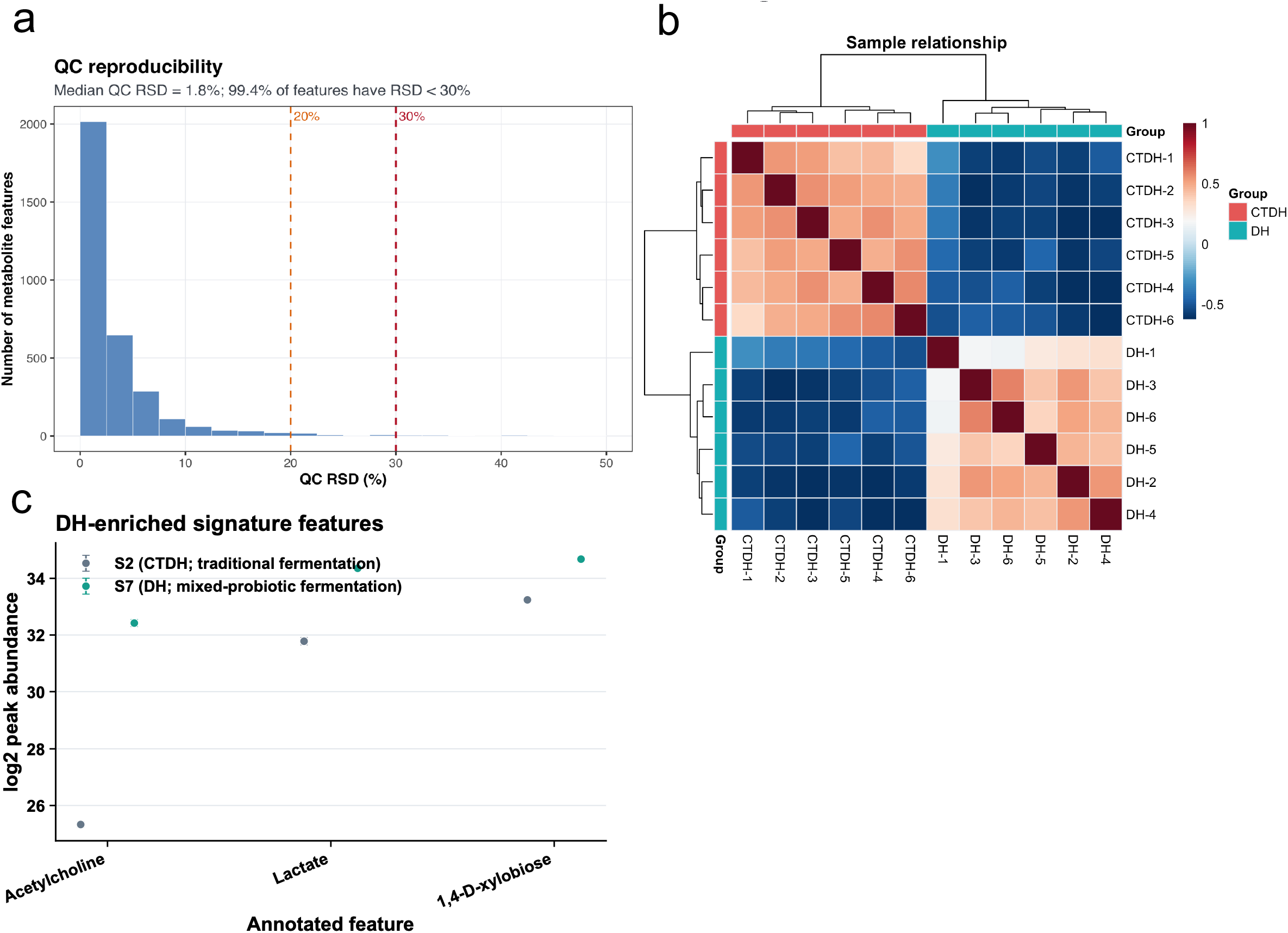
Quality control and relative signal-intensity profiles of selected metabolite features in untargeted metabolomics. (a) Frequency distribution of peak RSDs in QC samples (median RSD = 1.8%; 99.4% of features with RSD < 30%). (b) Pearson correlation matrix and hierarchical clustering among individual samples (CTDH-1–6 and DH-1–6). (c) Distinctive enrichment of signature features (Acetylcholine, Lactate, and 1,4-D-xylobiose) in the DH group.

**Supplementary Fig. S4.**
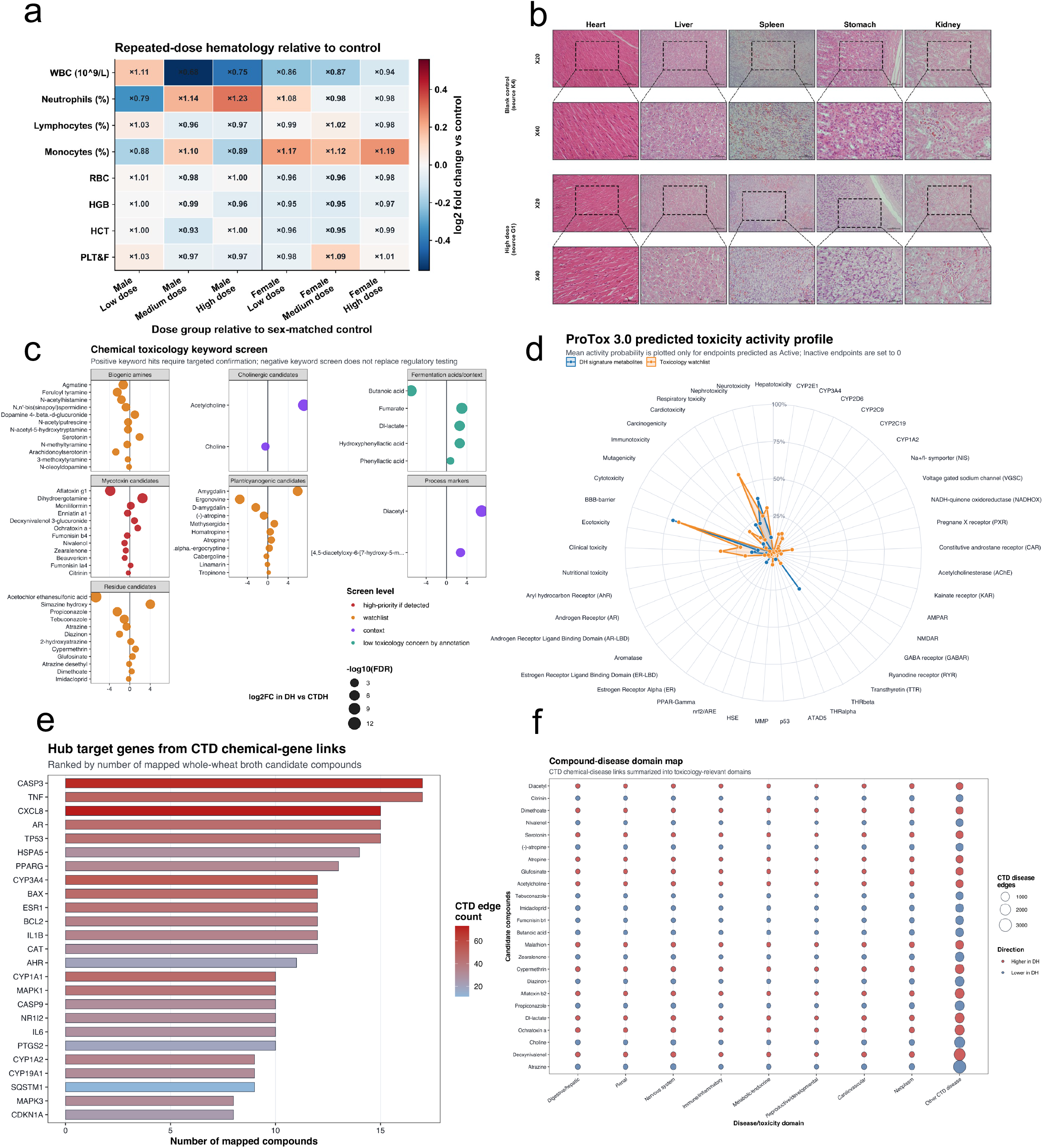
Hematological profiles, histopathology, and in silico toxicological assessment of putatively annotated metabolite features. (a) Repeated-dose hematological fold changes relative to sex-matched controls; color indicates log2 fold change. Overall differences were tested separately by sex using Kruskal–Wallis tests with Benjamini–Hochberg correction. (b) Representative H&E-stained heart, liver, spleen, stomach, and kidney sections from male and female blank-control and high-dose animals at ×20 and ×40 magnification. (c) Keyword-based toxicological watchlist of putatively annotated features. Position indicates S7/DH-versus-S2/CTDH log2 fold change, point size indicates −log10(FDR), and color indicates screening level. (d) ProTox 3.0 mean predicted activity probabilities for S7/DH signature and toxicology-watchlist features. (e) Human CTD hub genes ranked by the number of mapped candidate compounds; color indicates CTD interaction count. (f)CTD compound–disease associations summarized into nine disease domains. Point size indicates association count and color indicates the direction in S7/DH versus S2/CTDH. Panels c–f are hypothesis-generating and do not confirm compound identity or toxicity.

**Supplementary Fig. S5.**
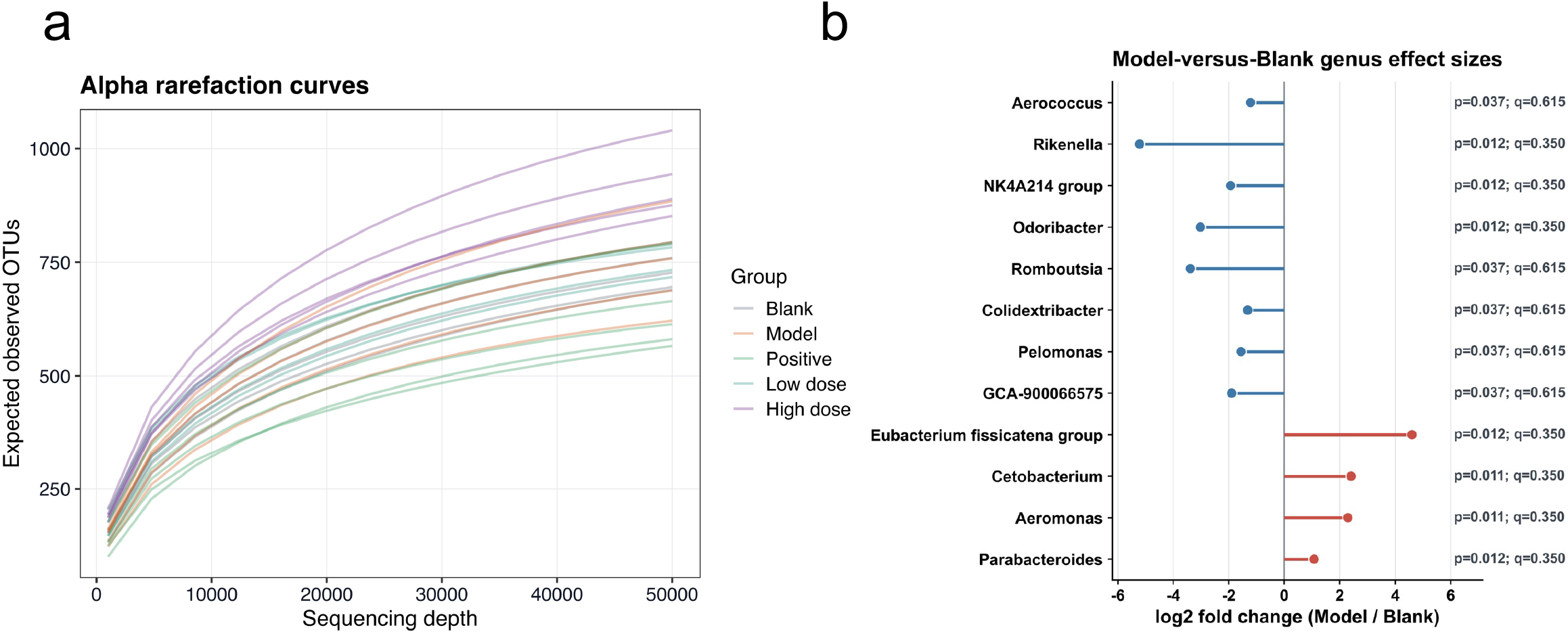
Sequencing-depth validation and effect-size analysis of model-associated genera. **(a)** Alpha rarefaction curves showing observed OTUs as a function of sequencing depth per sample, confirming sufficient coverage across groups. **(b)** Forest plot depicting log_2_ fold change of nominal model-responsive genera between Model and Blank groups (p < 0.05). (Note: q-values represent FDR-adjusted p-values; nominal effects are hypothesis-generating as individual taxa did not pass stringent multi-testing correction at FDR < 0.05).

**Supplementary Table S1.** Materials, reagents, and standards used in this study.

| No. | Category | Material/Reagent | Specification/Purity | Batch No. | Supplier/Source | Use/Note |
| --- | --- | --- | --- | --- | --- | --- |
| 1 | Enzyme | Cellulase | - | C8270 | Beijing Solarbio Science & Technology Co., Ltd., Beijing, China | Enzymatic hydrolysis |
| 2 | Enzyme | Pectinase | - | P8181 | Beijing Solarbio Science & Technology Co., Ltd., Beijing, China | Enzymatic hydrolysis |
| 3 | Reagent | Absolute ethanol | Analytical grade | 20220621 | Tianjin Tianli Chemical Reagent Co., Ltd., Tianjin, China | Extraction/analysis |
| 4 | Standard | Rutin reference standard | Reference standard | - | Beijing Solarbio Science & Technology Co., Ltd., Beijing, China | Total flavonoid assay |
| 5 | Standard | Gallic acid reference standard | Reference standard | - | Beijing Solarbio Science & Technology Co., Ltd., Beijing, China | Total polyphenol assay |
| 6 | Reagent | Folin-Ciocalteu reagent | - | 20230305 | Beijing Solarbio Science & Technology Co., Ltd., Beijing, China | Total polyphenol assay |
| 7 | Reagent | Sodium nitrite | Analytical grade | 20200918 | Tianjin Tianli Chemical Reagent Co., Ltd., Tianjin, China | Total flavonoid assay |
| 8 | Reagent | Aluminum nitrate | Analytical grade | 20230301 | Tianjin Damao Chemical Reagent Factory, Tianjin, China | Total flavonoid assay |
| 9 | Reagent | Sodium hydroxide | Analytical grade | 20221005 | Tianjin Damao Chemical Reagent Factory, Tianjin, China | pH adjustment/assay reagent |
| 10 | Reagent | Unspecified reagent | - | 20230305 | Beijing Solarbio Science & Technology Co., Ltd., Beijing, China | Reagent name to be confirmed |
| 11 | Reagent | Anhydrous sodium carbonate | Analytical grade | 20200801 | Tianjin Damao Chemical Reagent Factory, Tianjin, China | Total polyphenol assay |
| 12 | Solvent | Ultrapure water | - | - | Laboratory-prepared | General solvent |
| 13 | Raw material | Hetao wheat | Food material | - | Bayannur, Inner Mongolia, China | Whole-wheat substrate |
| 14 | Standard | D(+)-anhydrous glucose / D(+)-Glucose standard | CAS No. 50-99-7 | K8348399 | Shanghai Yuanye Bio-Technology Co., Ltd., Shanghai, China | Polysaccharide/glucose calibration |
| 15 | Buffer | pH 4.0 buffer solution | - | 20260411 | Tianjin Huasheng Chemical Reagent Co., Ltd., Tianjin, China | pH meter calibration |
| 16 | Buffer | pH 7.0 buffer solution | - | 20260117 | Tianjin Zhongtian Chemical Co., Ltd., Tianjin, China | pH meter calibration |
| 17 | Reagent | Phenol solution | 5% | 20260206 | Tianjin Zhongtian Chemical Co., Ltd., Tianjin, China | Phenol-sulfuric acid assay |
| 18 | Indicator | Bromocresol green-methyl red indicator | - | 20260411 | Tianjin Huasheng Chemical Reagent Co., Ltd., Tianjin, China | Nitrogen assay/titration indicator |
| 19 | Buffer | MES-TRIS buffer | 0.05 mol/L | 3330715 | Isekyu Biotechnology Co., Ltd. | Soluble dietary fiber assay |
| 20 | Enzyme | alpha-Amylase | 10000 U/mL ± 1000 U/mL | 260401 | Shanghai Lanji Technology Development Co., Ltd., Shanghai, China | Soluble dietary fiber assay |
| 21 | Enzyme | Amyloglucosidase | 2000-3300 U/mL | 260401 | Shanghai Lanji Technology Development Co., Ltd., Shanghai, China | Soluble dietary fiber assay |
| 22 | Enzyme | Protease solution | 300-400 U/mL | 260401 | Shanghai Lanji Technology Development Co., Ltd., Shanghai, China | Soluble dietary fiber assay |
| 23 | Reagent | Ethanol | 95%, analytical grade | 20260103 | Tianjin Damao Chemical Reagent Partnership, Tianjin, China | Soluble dietary fiber assay |
| 24 | Reagent | Ethanol | 78%, prepared from 95% ethanol | - | Laboratory-prepared | Soluble dietary fiber assay |
| 25 | Reagent | Acetone | Analytical grade | 2023121801 | Chengdu Kelong Chemical Co., Ltd., Chengdu, China | Soluble dietary fiber assay |
| 26 | Indicator | Phenolphthalein | Analytical grade | 20250214 | Tianjin Damao Chemical Reagent Factory, Tianjin, China | Titration indicator |
| 27 | Reagent | Boric acid | Analytical grade | 20190815 | Tianjin Damao Chemical Reagent Factory, Tianjin, China | Nitrogen assay |
| 28 | Reagent | Copper sulfate | Analytical grade | 20130906 | Tianjin Tianli Chemical Reagent Co., Ltd., Tianjin, China | Kjeldahl digestion catalyst |
| 29 | Reagent | Sodium hydroxide | Analytical grade | 20260102 | Tianjin Deen Chemical Reagent Co., Ltd., Tianjin, China | Nitrogen assay / NaOH solution preparation |
| 30 | Reagent | Potassium sulfate | Analytical grade | 201612011 | Tianjin Tianli Chemical Reagent Co., Ltd., Tianjin, China | Kjeldahl digestion catalyst |
| 31 | Reagent | Concentrated hydrochloric acid | Analytical grade | 2023103101 | Chengdu Kelong Chemical Co., Ltd., Chengdu, China | HCl solution preparation |
| 32 | Reagent | Concentrated sulfuric acid | Analytical grade | 2101297 | Xilong Scientific Co., Ltd., China | Digestion / phenol-sulfuric acid assay |

**Supplementary Tables S2-S4.**

|  |  |  |  |  |  |  |  |  |  |  |
| --- | --- | --- | --- | --- | --- | --- | --- | --- | --- | --- |
| **a** |  |  |  |  |  | **b** |  |  |  |  |
| Table S2. Factor levels for orthogonal design |  |  |  |  |  | Table S3. Orthogonal design and results |  |  |  |  |
| Level | A<br>Total enzyme dosage (%) | B<br>Hydrolysis temperature (°C) | C<br>Hydrolysis time (min) |  |  | Run | A | B | C | Composite score |
| 1 | 0.1 | 37 | 10 |  |  | 1 | 1 | 1 | 1 | 95.704 |
| 2 | 0.2 | 47 | 20 |  |  | 2 | 1 | 2 | 2 | 89.503 |
| 3 | 0.3 | 57 | 30 |  |  | 3 | 1 | 3 | 3 | 93.927 |
| **c** |  |  |  |  |  | 4 | 2 | 1 | 2 | 92.902 |
| Table S4. ANOVA for orthogonal design |  |  |  |  |  | 5 | 2 | 2 | 3 | 92.905 |
| Factor | Sum of squares | df | Mean square | F | Significance | 6 | 2 | 3 | 1 | 96.167 |
| A | 8.951 | 2 | 4.475 | 4.073 | * | 7 | 3 | 1 | 3 | 92.522 |
| B | 74.531 | 2 | 37.265 | 33.914 | ** | 8 | 3 | 2 | 1 | 90.533 |
| C | 13.723 | 2 | 6.862 | 6.244 | ** | 9 | 3 | 3 | 2 | 94.784 |
| Error | 21.976 | 20 | 1.099 |  |  | K1 | 279.134 | 281.128 | 282.404 |  |
|  |  |  |  |  |  | K2 | 281.974 | 272.941 | 277.189 |  |
|  |  |  |  |  |  | K3 | 277.839 | 284.878 | 279.354 |  |
|  |  |  |  |  |  | R | 4.108 | 11.937 | 5.215 |  |

**Supplementary Tables S5-S6.**

| Table S5. Factor levels for response surface design |  |  |  |
| --- | --- | --- | --- |
|  | A<br>Inoculum size (%) | B<br>Fermentation time (h) | C<br>Fermentation temperature (°C) |
| -1 | 10 | 20 | 27 |
| 0 | 12 | 24 | 32 |
| 1 | 14 | 28 | 37 |

b
| Table S6. Response surface design and results |  |  |  |  |
| --- | --- | --- | --- | --- |
| Run | A | B | C | Composite score Y |
| 1 | 10 | 20 | 32 | 72.0032 |
| 2 | 14 | 20 | 32 | 75.0165 |
| 3 | 10 | 28 | 32 | 78.7989 |
| 4 | 14 | 28 | 32 | 75.6941 |
| 5 | 10 | 24 | 27 | 67.6336 |
| 6 | 14 | 24 | 27 | 82.1484 |
| 7 | 10 | 24 | 37 | 70.2858 |
| 8 | 14 | 24 | 37 | 56.9974 |
| 9 | 12 | 20 | 27 | 81.2691 |
| 10 | 12 | 28 | 27 | 83.3692 |
| 11 | 12 | 20 | 37 | 71.8185 |
| 12 | 12 | 28 | 37 | 71.2671 |
| 13 | 12 | 24 | 32 | 85.0527 |
| 14 | 12 | 24 | 32 | 84.2066 |
| 15 | 12 | 24 | 32 | 83.1108 |
| 16 | 12 | 24 | 32 | 90.9837 |
| 17 | 12 | 24 | 32 | 85.0738 |

c
| Source | Sum of squares | df | Mean square | F value | p value |
| --- | --- | --- | --- | --- | --- |
| Model | 1082.31 | 9 | 120.26 | 19.86 | 0.0003 |
| A-Inoculum size | 0.1610 | 1 | 0.1610 | 0.0266 | 0.8751 |
| B-Fermentation time | 10.17 | 1 | 10.17 | 1.68 | 0.2359 |
| C-Fermentation temperature | 242.57 | 1 | 242.57 | 40.07 | 0.0004 |
| AB | 9.36 | 1 | 9.36 | 1.55 | 0.2538 |
| AC | 193.26 | 1 | 193.26 | 31.92 | 0.0008 |
| BC | 1.76 | 1 | 1.76 | 0.2903 | 0.6067 |
| A<sup>2</sup> | 339.99 | 1 | 339.99 | 56.16 | 0.0001 |
| B<sup>2</sup> | 7.35 | 1 | 7.35 | 1.21 | 0.3069 |
| C<sup>2</sup> | 232.64 | 1 | 232.64 | 38.43 | 0.0004 |
| Residual | 42.38 | 7 | 6.05 |  |  |
| Lack of fit | 4.72 | 3 | 1.57 | 0.1670 | 0.9134 |
| Pure error | 37.66 | 4 | 9.42 |  |  |
| Corrected total sum of squares | 1124.69 | 16 |  |  |  |
| R<sup>2</sup> | 0.9623 |  |  |  |  |
| Adjusted R<sup>2</sup> | 0.9139 |  |  |  |  |

Panel definitions: (a) Factor levels for the Box–Behnken response-surface design. (b) Box–Behnken experimental design and corresponding composite scores. (c) ANOVA and model-fit statistics for the quadratic response-surface model.

